# Projectome-defined subtypes and modular intra-hypothalamic subnetworks of peptidergic neurons

**DOI:** 10.1101/2023.05.25.542241

**Authors:** Zhuolei Jiao, Taosha Gao, Xiaofei Wang, Wen Zhang, Nasim Biglari, Emma E Boxer, Lukas Steuernagel, Xiaojing Ding, Zixian Yu, Mingjuan Li, Mingkun Hao, Hua Zhou, Xuanzi Cao, Shuaishuai Li, Tao Jiang, Jiamei Qi, Xueyan Jia, Zhao Feng, Biyu Ren, Yu Chen, Xiaoxue Shi, Dan Wang, Xinran Wang, Luyao Han, Yikai Liang, Congcong Wang, E Li, Yue Hu, Zi Tao, Humingzhu Li, Xiang Yu, Min Xu, Hung-Chun Chang, Yifeng Zhang, Huatai Xu, Jun Yan, Anan Li, Qingming Luo, Ron Stoop, Scott M. Sternson, Jens C. Brüning, David J. Anderson, Mu-ming Poo, Hui Gong, Yangang Sun, Xiaohong Xu

## Abstract

The hypothalamus plays a vital role in coordinating essential neuroendocrine, autonomic, and somatomotor responses for survival and reproduction. While previous studies have explored population-level projections of hypothalamic neurons, the specific innervation patterns of individual hypothalamic axons remain unclear. To understand the organization of hypothalamic axon projections, we conducted a comprehensive reconstruction of single-cell projectomes from 7,180 mouse hypothalamic neurons expressing specific neuropeptides. Our analysis identified 31 distinct subtypes based on projectome-defined characteristics, with many exhibiting long-range axon collateral projections to multiple brain regions. Notably, these subtypes selectively targeted specific subdomains within downstream areas, either unilaterally or bilaterally. Furthermore, we observed that individual peptidergic neuronal types encompassed multiple projectome-defined subtypes, explaining their diverse functional roles. Additionally, by examining intra-hypothalamic axon projections, we uncovered six modular subnetworks characterized by enriched intramodular connections and distinct preferences for downstream targets. This modular organization of the intra-hypothalamic network likely contributes to the coordinated organization of hypothalamic outputs. In summary, our comprehensive projectome analysis reveals the organizational principles governing hypothalamic axon projections, providing a framework for understanding the neural circuit mechanisms underlying the diverse and coordinated functions of the hypothalamus.

## Main text

The hypothalamus is an evolutionarily ancient brain structure at the ventral base of the brain, comprising at least 40 tightly packed nuclei and millions of intricately connected neurons^1–4^. It regulates neuroendocrine, autonomic, and somatomotor responses in a coordinated manner for essential behaviors and maintains homeostasis of many physiological processes. For example, the hypothalamus controls body temperature, metabolism, hormone secretion, eating, drinking, fluid balance, sleep-wake cycle, circadian rhythm, stress responses, predator defense, reproductive behaviors, and reward processing. Yet, despite the well-recognized importance of the hypothalamus, the organization of hypothalamic axon projections that underlies its coordinated regulation of this plethora of physiological and behavioral functions remains to be fully understood.

By injecting an anterograde tracer such as Phaseolus vulgaris leucoagglutinin into specific rat hypothalamic nuclei, previous studies have revealed extensive hypothalamic projections throughout the brain and intricated reciprocal intra- hypothalamic connections^5–12^. These macroscopic studies have illuminated the overall axon projections of some hypothalamic nuclei, but axon projections of molecularly defined neuronal subtypes within the same nucleus could not be differentiated. Furthermore, an overview of the axon organization of the entire hypothalamus remains to be explored.

More recently, single-cell RNA-seq (scRNA-seq) analyses were used to identify neuronal types in the mouse hypothalamus based on gene expression patterns^13–15^. Many transcriptome-defined hypothalamic neuron types express genes encoding specific neuropeptides such as *agouti-related peptide* (*Agrp*), *proopiomelanocortin (Pomc), oxytocin (Oxt), and Orexin*, which are evolutionarily conserved neuromodulators of essential functions^16^. Additionally, virus-mediated anterograde tracing has delineated projections of these neuropeptide-expressing neurons at a population level^17–19^. However, local activation or inhibition of neuronal populations expressing a single neuropeptide often affects multiple behaviors and physiological processes^20, 21^. This suggests that transcriptionally similar neurons may exert distinct functions by innervating separate sets of downstream targets via axon collaterals, thereby regulating multiple processes in a coordinated manner. In other words, distinct projection-based subtypes may exist within a single transcriptome-defined neuron type to promote functional diversity. Thus, delineating the whole-brain projection pattern of single axons (projectomes) is pivotal for understanding neural mechanisms underlying hypothalamic functions.

In this study, we used a recently developed axon tracing method based on fluorescence micro-optical sectioning tomography (fMOST)^22^ and constructed the single-cell projectomes of ∼7000 hypothalamic neurons that express specific neuropeptide genes. This provides the largest known single-neuron projectome dataset for the hypothalamus. We further identified 31 projectome-defined subtypes, many of which sent projections to specific ipsilateral or bilateral subdomains of targeted regions via their axon collaterals. Notably, each neuropeptide-defined population comprised multiple projectome-defined subtypes, providing a structural basis for its diverse but coordinated functions. Moreover, by analyzing intra-hypothalamic axon projection patterns, we identified six modular subnetworks with enriched recurrent intra-modular connections, each showing preferences for distinct target areas. Such a modular subnetwork organization may serve as an intra-hypothalamic coordination of physiological homeostasis and innate behaviors. Overall, this single-neuron projectome dataset reveals organization principles of hypothalamic axon projections and provides a valuable resource for future functional studies.

### Tracing single-cell projectomes of hypothalamus peptidergic neurons

To achieve bright, sparse, and specific labeling of molecularly defined hypothalamic neurons, we injected the hypothalamus with adeno-associated virus (AAVs) carrying Cre-dependent GFP in genetically modified mice that expressed *Cre* recombinase under the control of a neuropeptide promoter; alternatively, we injected AAVs that expressed GFP driven by an *Orexin* promotor in wild-type mice to label *Orexin*-expressing neurons (**Fig. 1a**, Extended Data Fig.1, Supplementary Table1, see Methods). After sufficient viral expression, brain samples were processed and imaged by fMOST, which could achieve a high spatial resolution (0.32μm * 0.32μm * 1μm) for the whole-brain tissue. We traced the projection of individually labeled axons by a streamlined semi-automatic procedure previously described^23^ (**Fig. 1a**, see Methods). To permit cross-sample comparison, we registered all traced projections onto the Allen mouse brain common coordinate frame (CCFv3)^24^ (Extended Data Fig.2a, Supplementary Table 2 for brain structure abbreviations). In total, we reconstructed the brain-wide projectome of 7180 individual neurons from 16 peptidergic populations whose somata were registered to the hypothalamus (**Fig. 1a-1b**).

**Fig. 1.**
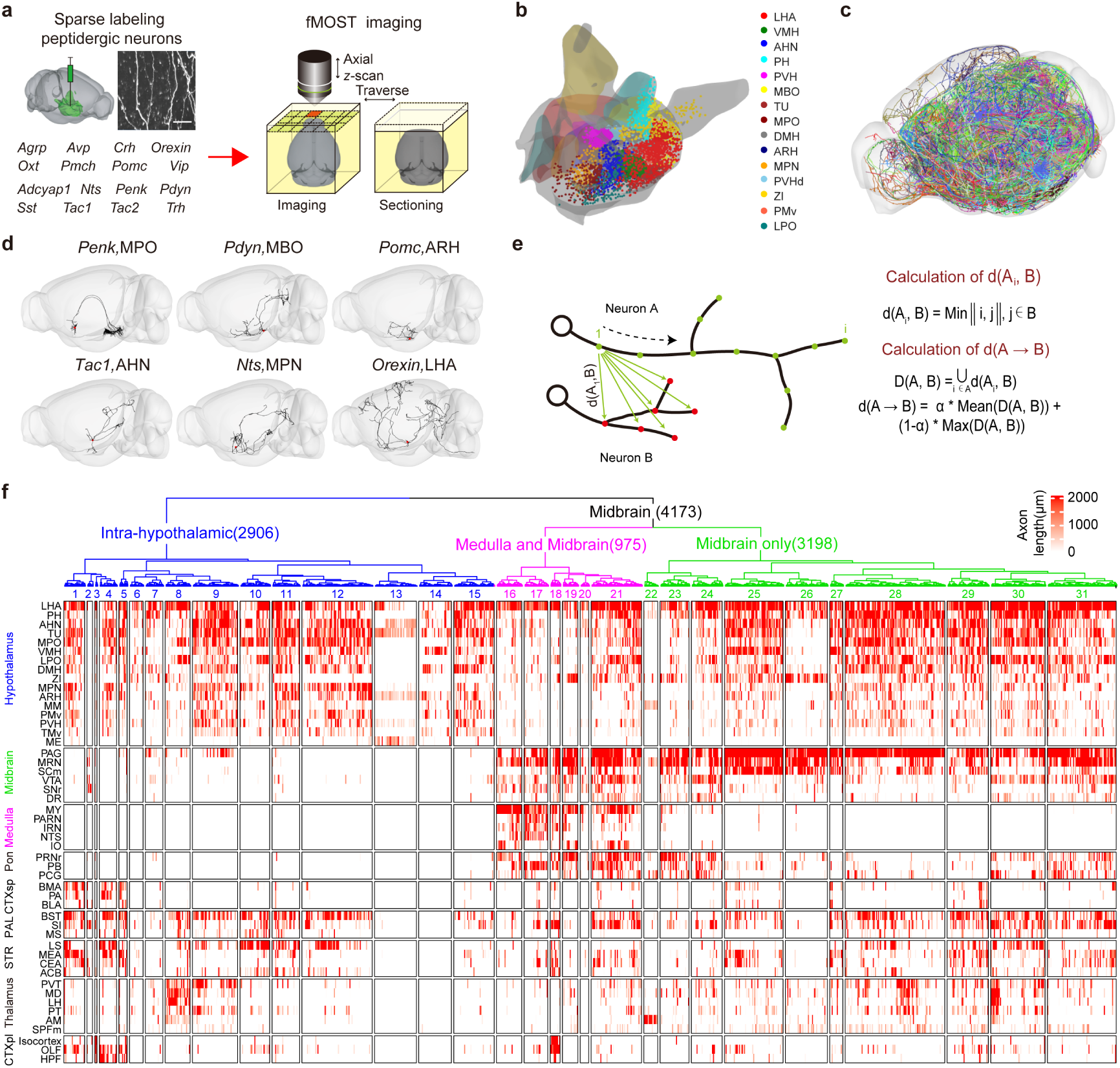
Single-cell projectomes of hypothalamic peptidergic neurons. **a.** The labeling and imaging procedure. We sparsely labeled hypothalamic neurons expressing one of the 16 indicated neuropeptides (left, bottom) and imaged the brain sample on an fMOST platform (right). The fluorescent image shows the processes of a labeled neuron. Scale bar, 50 μm. **b.** Soma distribution of the 7180 reconstructed peptidergic neurons in various hypothalamic nuclei (color-coded, see Supplementary Table 2 for nomenclature) in a 3D view. All neurons were mapped to the same hemisphere of the brain. **c.** Axon projections of all 7180 reconstructed neurons throughout the brain. Each neuron was labeled with a different color. **d.** Examples of the reconstructed morphology of individual neurons in a whole-brain view. The expressed neuropeptide and the soma location of the neuron were indicated above the brain. **e.** The algorithm for calculating the similarity between neuron pairs. Briefly, for each point in neuron A, we calculated the closest distance to points in neuron B, defined as dis (Ai), as shown on the left. For all points in neuron A, we calculate a mean and max value of all dis (Ai)s. We weighted these mean and max values by the portion of points in neuron A that had a dis (Ai) value below or above the mean and defined this as the directional A→B distance. The distance between the A&B neuron pair was the average of the two directional (A→B & B→A) distances. **f.** A summary of axon projection length (in μm) in each brain area labeled on the left (in rows) for the 31 projectome-defined subtypes, labeled #1-#31. We divided these subtypes into three major classes (color-coded, blue, intra-hypothalamic; pink, medulla & midbrain projecting; green, only midbrain projecting). Each tick represents the value of the axon projection length of a single neuron in a heatmap fashion with the scale shown above. *See also Extended Data Fig.1–4, Supplementary Table 1–3*.

The soma distribution of these reconstructed neurons closely matched the known expression patterns of these neuropeptides. Specifically, substantially more neurons were obtained for broadly expressed neuropeptides (*Adcyap1, Nts, Penk, Pdyn, Sst, Tac1, Tac2, Trh*) than those for more restrictedly expressed ones (*Agrp, Avp, Crh, Orexin, Oxt, Pmch, Pomc, Vip*) (Extended Data Fig.2b). Overall, reconstructed neurons occupied most hypothalamic nuclei, showing a whole-hypothalamus coverage of our dataset (**Fig. 1b**, Extended Data Fig.2c). Furthermore, these 16 selected neuropeptides served as molecular markers for about half of the 66 transcriptional-defined neuronal types in the mouse hypothalamus (Extended Data Fig.3)^13^, covering both excitatory and inhibitory neuronal types, and some neuropeptides such as *Orexin* and *Pomc* marking single transcriptome-defined neuron types. Moreover, the total axon arbors of these 7180 neurons covered ∼ 21.8% and ∼ 10.2% of the voxels (voxel size 25 μm) for the entire brain volume on the ipsilateral and contralateral side, respectively **(Fig. 1c)**. Thus, our single-neuron projectome dataset reasonably represents mouse hypothalamic neurons.

### Single-cell projectome-based classification of hypothalamic neurons

At first glance, individual hypothalamic neurons showed high variation in their axon arborization patterns and target areas (**Fig. 1d**), making classification based on simple projection target selectivity unrealistic. We thus calculated morphological similarity between neuronal pairs by computing the weighted average of the closest distances for all traced points of the reconstructed neuron (**Fig. 1e**, see Methods). Using hierarchical agglomerative grouping of the similarity matrix, we identified 31 projectome-defined neuronal subtypes, as shown in **Fig. 1f**. Moreover, each subtype encompassed multiple peptidergic populations (Supplementary Table 3).

These 31 subtypes could be roughly grouped into three major classes based on whether they projected to the midbrain and medulla: (1) Intra-hypothalamic class (subtypes #1 to #15); (2) medulla and midbrain class (subtypes #16 to #21); and (3) midbrain only class (subtypes #22 to #31). Neuronal projections to other brain areas (e.g., cortex, pallidum, thalamus, striatum, hippocampus, and olfactory area) did not show such group selectivity, although each projectome subtype had distinct preference for these targeted brain areas (**Fig. 1f**). The emergence of target selectivity among projectome-defined subtypes supports our classification method.

To examine whether the projectome-defined neuron subtypes/classes described above were specific to peptidergic neurons, we used the method described above (now published as a Python library (https://pypi.org/project/pyswcloader/) to analyze the projectome of 77 hypothalamic neurons in a previous dataset^25^ generated by using a pan-neuronal labeling method and light-sheet imaging. We found that these 77 hypothalamic neurons could be similarly grouped into the above three classes (Extended Data Fig.4). Thus, projectome-defined hypothalamic neuron classes and subtypes could be generalized to hypothalamic neurons reconstructed via other labeling and imaging methods. Taken together, we have identified three major classes and 31 projectome-defined subtypes of hypothalamic neurons.

### Characteristics of the intra-hypothalamic projecting class

Intra-hypothalamic projecting subtypes #1 to #15 innervated mainly hypothalamic nuclei and lacked long-range projections to midbrain regions (**Fig. 1f**). Nevertheless, neurons in nearly half of these subtypes made preferential long-range projections to non-midbrain areas, including the amygdala, cortex, hippocampus, and thalamus (**Fig. 2a**), providing part of the features that differentiated these subtypes among the intra-hypothalamic projecting class. For example, subtypes #1 and #5 neurons showed strong projections to the central amygdala nucleus (CEA) and medial amygdala nucleus (MEA) and modest projections to the posterior amygdala nucleus (PA) (**Fig. 2b**). By comparison, subtype #4 neurons showed modest projections to amygdala nuclei and prominent projections to CA1 of the hippocampus (**Fig. 2c**). Meanwhile, subtype #2 specifically projected to the substantia nigra (SNr) (**Fig. 2d**), whereas subtype #3 projected to both the anterior cingulate area (ACA) and somatomotor areas (MO) (**Fig. 2e**). Subtype #8 and #9 neurons projected to thalamic nuclei, with disparate preferences to lateral habenula (LH), mediodorsal nucleus of the thalamus (MD), and paraventricular nucleus of the thalamus (PVT) (**Fig. 2f**).

**Fig. 2.**
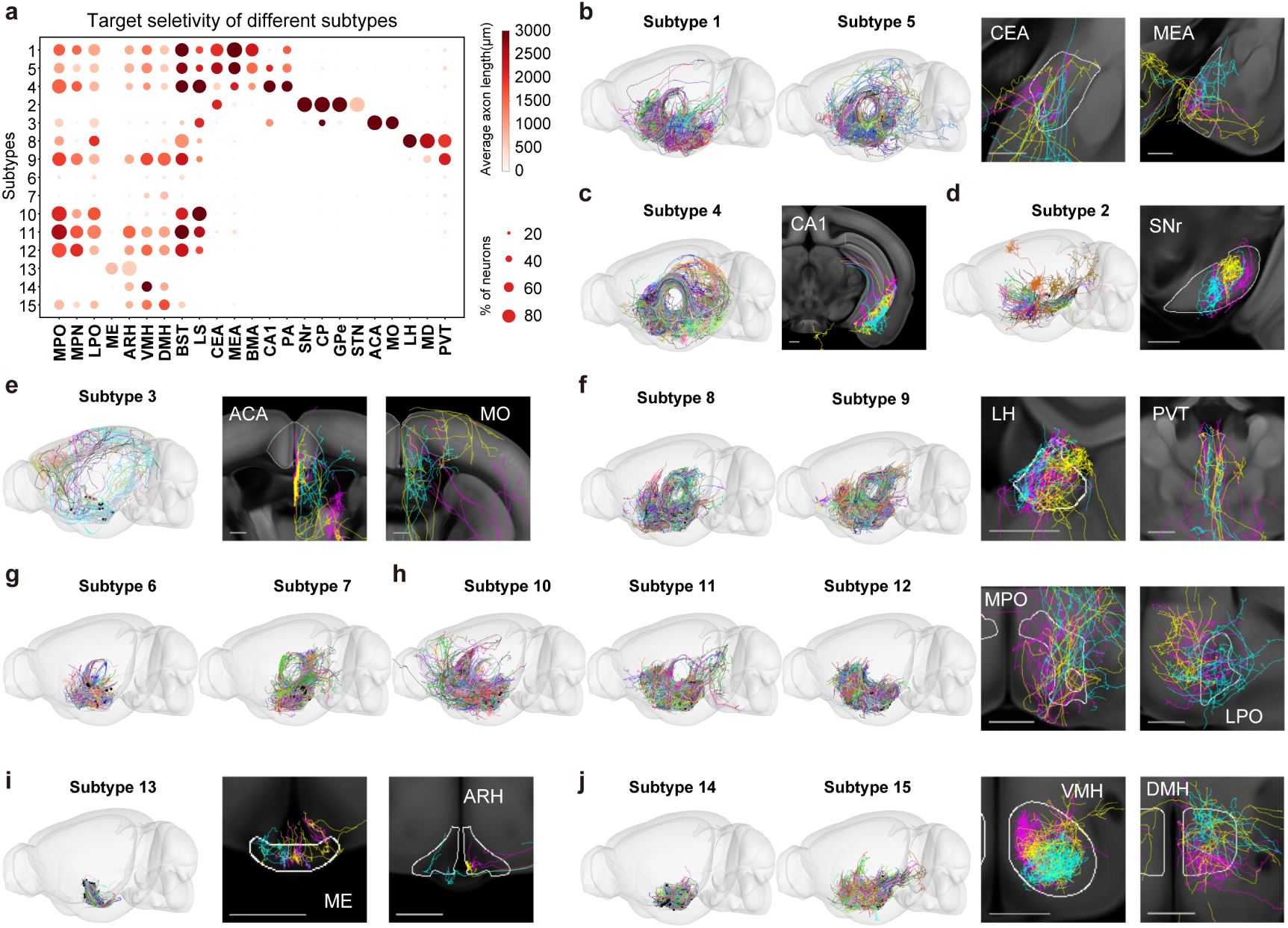
Characteristics of the intra-hypothalamic projecting class. **a.** A dot plot depicting selective projections of each subtype (row) in each brain area (column). The dot circle’s size and color intensity indicate the percentage of neurons in each subtype that projected to the indicated brain area and the average projection length (in μm) in a heatmap fashion, respectively, with the scale on the right. **b-j.** The total axon projections of neurons of the indicated subtype are plotted in a 3D brain on the left. Zoom-in views of representative axon projections from three representative neurons (color-coded) from each subtype in defined subdomains of the indicated target areas are shown on the right. Scale bar, 500 μm.

Axons of the rest of subtypes of the intra-hypothalamic projecting class more frequently terminated within and around the hypothalamus. The subtypes #6 and #7 neurons projected only locally around the soma, whereas the subtypes #10 to #15 neurons projected more extensively to other hypothalamic nuclei such as medial preoptic area (MPO), lateral preoptic area (LPO), ventral medial hypothalamic nucleus (VMH), and dorsal medial nucleus of the hypothalamus (DMH) (**Fig. 2g-j**), as well as to bed nuclei of stria terminalisis (BST) and lateral septum (LS) outside of the hypothalamus. An exception was subtype #13 neurons, which showed restricted projections to the median eminence (ME) and arcuate nucleus (ARH) (**Fig. 2i**). In line with previous studies on neurohypophysis^26, 27^, the soma of subtype #13 neurons was located primarily in PVH, and 180 out of the 306 subtype #13 neurons expressed *Oxt* (Supplementary Table 3).

### Characteristics of the midbrain projecting classes

As shown in **Fig. 1f**, subtypes #16 to #31 differed significantly from #1 to #15 by their projections to midbrain areas. Interestingly, within the hypothalamus, these midbrain-projecting neurons also showed higher projection strength (as defined by the total length of axon arbors covering the targeted region) in the lateral hypothalamic area (LHA), posterior hypothalamus nucleus (PH), and zona incerta (ZI) and lower strength in the medial preoptic nucleus (MPN), ARH, ventral premammillary nucleus (PMv), and the ventral part of the tuberomammillary nucleus (TMv) (Extended Data Fig.5a), as compared to non-midbrain-projecting neurons. Among all midbrain projecting neurons, 23% had collaterals projecting to the medulla (subtypes #16 to #21) (**Fig. 1f**), most of which projected strongly to motor-related areas in the medulla (Extended Data Fig.5b). Some also innervated distinct unilateral and bilateral non-motor targets of the medulla and pons areas (Extended Data Fig.5c–d). Thus, individual axons emanating from the medulla-projecting neurons arborize extensively, and their collateral branches could innervate multiple regions in the midbrain, pons, and medulla (Extended Data Fig.5e).

Among the midbrain areas receiving hypothalamic projections, the most prominent was the periaqueductal gray (PAG) (Extended Data Fig.6a), which includes anterior PAG (aPAG) and rostral nuclei (INC, Su3, ND, and PRC), as well as four longitudinally organized PAG columns (Paxino) - dorsomedial (dm), dorsolateral (dl), lateral (l), and ventrolateral (vl) PAG (**Fig. 3a**). Among all PAG-projecting neurons, 74.4% and 12.4 % exhibited ipsilateral and contralateral preferences, respectively. Moreover, neurons of different subtypes showed distinct projection preferences (**Fig. 3b-d**). For example, subtypes #23 and #31 mainly projected ipsilaterally, whereas #27 slightly preferred the contralateral side (**Fig. 3c-d**). We next determined the column/nucleus preferences of PAG-projecting neurons on the ipsilateral side by calculating the fold difference of projection density (total arbor length divided by the volume of targeted area) for a particular column/nucleus relative to the average projection density for all areas. With the criterion of 3-fold difference, we found that PAG-projecting neurons that preferred distinct PAG regions (Extended Data Fig.6b). Column/nucleus preferences also varied among various subtypes (**Fig. 3e**). Furthermore, the column/nucleus preferences of bilateral PAG-projecting neurons exhibited striking similarity between the two hemispheres (**Fig. 3f**), providing a structural basis for bilateral control of PAG functions by a single bifurcating hypothalamic axon.

**Fig. 3.**
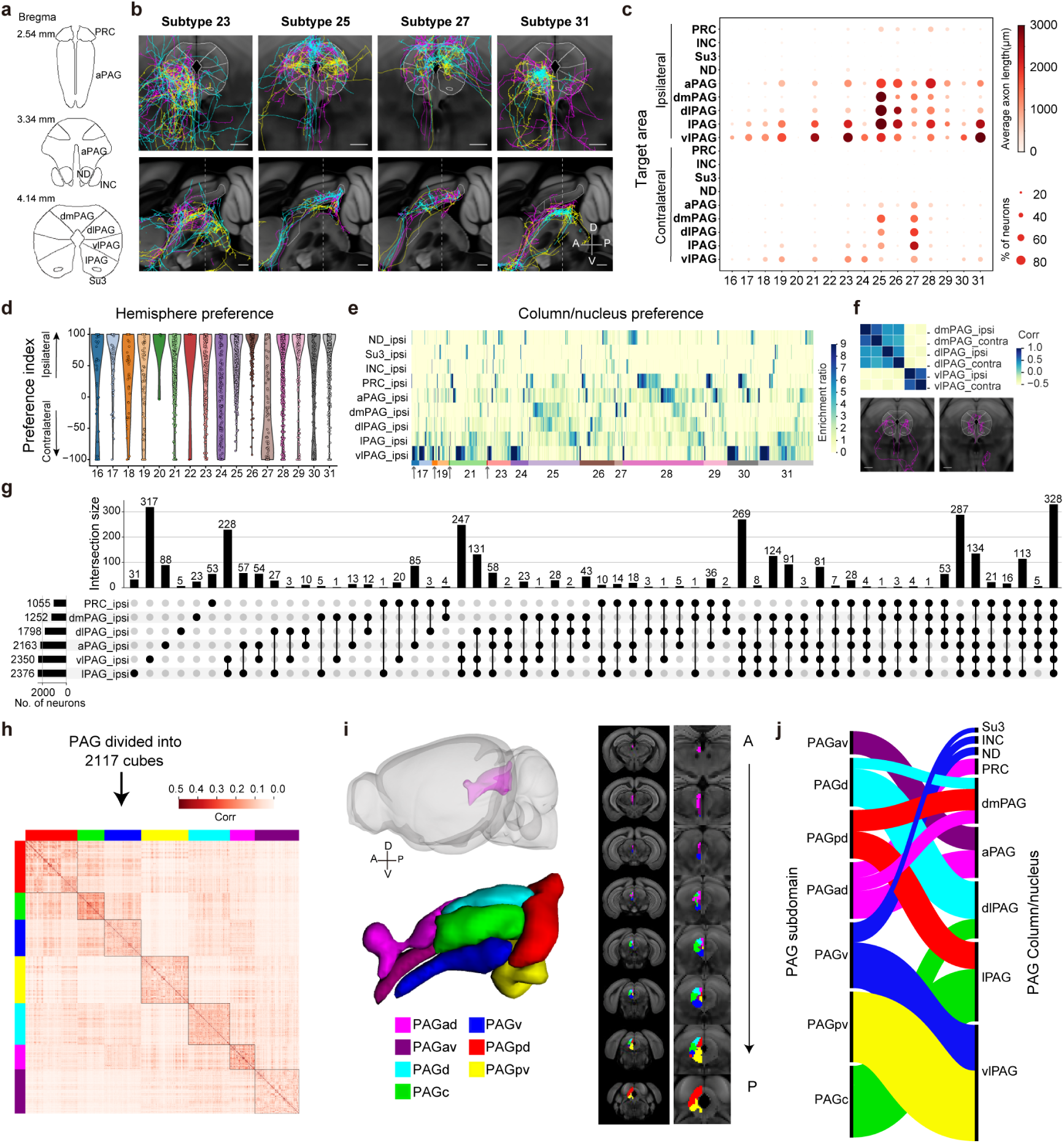
Complex and correlated axon patterns of midbrain projecting hypothalamic neurons in PAG. **a.** Schematics showing the nuclei and column structures of the PAG, which includes anterior PAG (aPAG) and rostral nuclei (INC, Su3, ND, and PRC), as well as four longitudinally organized PAG columns - dorsomedial (dm), dorsolateral (dl), lateral (l), and ventrolateral (vl) PAG **b.** Zoom-in views of representative axon projections of three neurons (color-coded) from the indicated subtype in a coronal (top) and sagittal (bottom) plane of a PAG section. Scale bar, 500 μm. Dashed lines showed the position of the coronal section on the sagittal plane. **c.** A dot plot depicting selective projections of each subtype (row) in each PAG column/nucleus on the ipsilateral or contralateral side. The dot circle’s size and color intensity indicate the percentage of neurons in each subtype that projected to the indicated brain area and the average projection length (in μm) in each column/nucleus in a heatmap fashion with the scale shown on the right. **d.** Violin plots of the preference index for the ipsilateral or contralateral PAG, calculated as the differences between the hemisphere divided by the sum of both hemispheres for each neuron in each subtype, represented by individual circles. **e.** Heatmap representation (sorted) of column/nucleus preference score for projections in the ipsilateral PAG, calculated as the fold difference of projection density (total arbor length divided by the volume of targeted area) for a particular column/nucleus relative to the average projection density for all areas, of individual neurons in each subtype. **f.** Correlation (corr) analysis of preference index in the ipsilateral (ipsi) and contralateral side (contra) of the indicated PAG column/nucleus for bilaterally PAG-projecting neurons. Representative images on the bottom show a coordinated innervation pattern of similar column/nucleus in both hemispheres by two bilaterally PAG-projecting neurons. **g.** An upset plot showing the intersection size of PAG-projecting neurons targeting each PAG column/nucleus and combinations of different PAG subdomains. Most PAG-projecting neurons target multiple column/nucleus. **h-j.** New PAG subdomains defined by co-innervation patterns of hypothalamic axons. **h.** We divided the PAG into 2117 100 μm-size cubes and calculated the pair-wise inter-cube correlation (corr) of hypothalamic projections. This way, seven modules (color-coded) showed higher intra-module than the inter-module correlation of hypothalamic axon projections. **i.** These seven subdomains were plotted in a 3D (left) and a 2D view (right) of the PAG. ad, anterio-dorsal; av, anterio-ventral; d, dorsal; c, central; v, ventral; pd, posterio-dorsal; pv, posterio-ventral. **j.** Schematics showing spatial correspondence between the newly defined PAG subdomains with atlas-defined PAG columns/nuclei. These innervation-based PAG subdomains rearranged the previously defined PAG columns and nuclei, with the overall dorsoventral and anterio-posterior patterns preserved. *See also Extended Data Fig.5–6*.

Taking advantage of the single-cell resolution of our dataset, we further examined projection patterns of individual PAG-projecting neurons regardless of their subtypes, focusing only on the ipsilateral side. We discovered some intriguing patterns. First, most PAG-projecting neurons targeted multiple columns/nuclei (**Fig. 3g**). Second, the pattern of PAG co-innervation was non-random, with vlPAG/lPAG pairing and dmPAG/dlPAG pairing frequently observed (**Fig. 3g**). Such co-innervation of different PAG columns/nuclei implies the co-regulation of PAG neurons across anatomically defined boundaries. To re-define PAG subdomains based on hypothalamic co-innervation patterns, we subdivided PAG into 100 μm-size cubes, calculated the pair-wise correlation of hypothalamic axon arbors within each cube, and performed clustering analysis of the correlation matrix (**Fig. 3h**). This yielded seven PAG subdomains, each exhibiting highly correlated hypothalamic projections (**Fig. 3i**). These innervation-based PAG subdomains rearranged the previously defined PAG columns and nuclei, with the overall dorsoventral and anterio-posterior patterns preserved (**Fig. 3i-j**). Interestingly, based on the published singe-cell projectomes of prefrontal cortex neurons^23^, a similar correlation analysis identified PAG subdomains that closely matched those defined by hypothalamic singe-cell projectomes (Extended Data Fig.6c), supporting the notion that these projectome-defined PAG subdomains might be functionally relevant. Thus, single-cell projectomes have revealed complex and patterned arborization of midbrain-projecting hypothalamic neurons within distinct PAG subdomains.

### Multiple projectome subtypes of Orexin-expressing neurons

Having categorized hypothalamic neurons based on single-cell projectomes, we further inquired whether there are distinct projectome-defined subtypes within a transcriptome-defined neuron type, which could offer a structural explanation for its functional diversity. To examine this question, we first focused on *Orexin*-expressing neurons, a population often characterized as a single transcriptionally defined excitatory neuron type by scRNA-seq analysis (Extended Data Fig.3) but known to exert complex regulatory functions on behaviors^28^. Previous studies have described axon projections of *Orexin* neurons at the population level^17^, indicating four projection routes: the ventral or dorsal ascending and descending pathways (**Fig. 4a**). However, whether individual *Orexin* axons project through one or more paths and their targeting patterns was unclear.

**Fig. 4.**
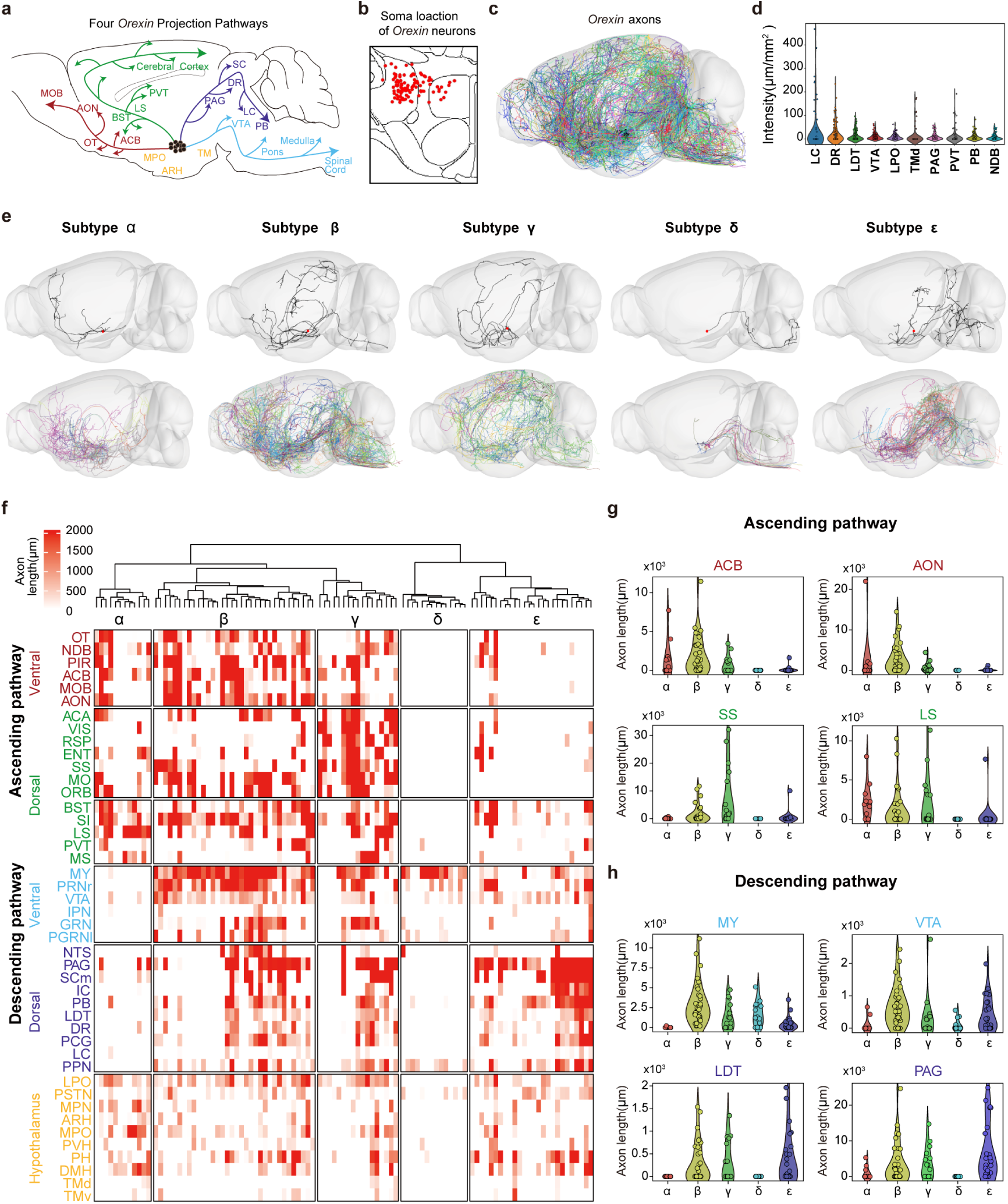
Multiple projectome subtypes of *Orexin*-expressing neurons. **a.** Schematics depicting four axon projection pathways (color-coded) of *Orexin* neurons described in previous studies: the ventral ascending (red), dorsal ascending (green), ventral descending (blue), and dorsal descending pathways (purple). **b.** The soma location of the 102 *Orexin* neurons is shown in a coronal section. **c.** Axon projections of the 102 *Orexin* neurons (color-coded) throughout the brain. **d.** Axon projections intensity of the 102 *Orexin* neurons in indicated brain areas. Each circle represents an individual neuron. **e.** Five projectome-defined subtypes of *Orexin* neurons, termed “*α*” through “*ε*”, were identified via morphology-based clustering analysis. Projections of a representative neuron (up panels) and all neurons (lower panels) for each subtype were plotted. **f.** A summary of axon projection length (in μm) for individual neurons of the five projectome-defined subtypes in each brain area labeled on the left. Each tick represents the value of the axon projection length of a given neuron in a heatmap fashion with the scale shown above. **g-h.** Violin plot of the projection length (in μm) of individual neurons in the five projectome-defined subtypes in specific target areas along the ascending (g) or descending (h) pathways. Each circle represents an individual neuron. *See also Extended Data Fig.7*

We here analyzed single-cell projectomes of all 102 *Orexin* neurons in our dataset and found their somata were primarily localized to LHA, as expected (**Fig. 4b**). Their axon projections extended broadly throughout the brain, innervating many downstream targets in the forebrain, thalamus, and midbrain structures (**Fig. 4c-d**). Notably, individual *Orexin* neurons exhibited large variability in their morphology and axon projection patterns (**Fig. 4e**). Morphology-based clustering analysis (see Fig. 1) of these 102 *Orexin* neurons yielded five projectome-defined subtypes, named “*α*” through “*ε*” (**Fig. 4e**). Although the somata of these projectome subtypes were intermingled (Extended Data Fig.7a), their axons showed systematic differences among projections along the four pathways described above. Specifically, subtype *α* mainly projected through the ascending pathways, whereas subtype *δ* and *ε* projected via descending pathways; subtype *β* and *γ* projected through both ascending and descending pathways. Consistently, subtypes *α, β,* and *γ* showed strong innervation of the cortex and forebrain structures. Subtypes *δ* and *ε* had very few cortical targets with *δ* subtype projecting solely posteriorly to the medulla (MY) and *ε* subtype showing strong projections mainly to PAG and pons (**Fig. 4e-h**). Differences also existed among the subtypes *α, β, and γ* in their cortical targets. The extensiveness of their cortical arborizations could be roughly ordered as *γ > β > α* (**Fig. 4f**, Extended Data Fig.7b). For example, the visual areas (VIS) and the retrosplenial area (RSP) were preferentially innervated by subtype *γ* (Extended Data Fig.7b). Correlation analysis of the projection strength of individual neurons further revealed coordinated targeting of cortical domains by individual *Orexin* neurons (Extended Data Fig.7c). Thus, *Orexin* neurons comprise multiple projectome subtypes of distinct axon innervation patterns. Such organization of projection patterns may provide a structural basis for the diverse functions of *Orexin* neurons.

### Distinct projectome subtypes with arcuate Agrp and Pomc neurons

We next compared the single-cell projectomes of *Agrp* (n = 62) and *Pomc* (n = 129) neurons located within the arcuate nucleus of the hypothalamus in the dataset (**Fig.5a**). These two transcriptome-defined neuron types (Extended Data Fig.3) exert antagonistic actions on the feeding behavior: activation of *Agrp* and *Pomc* neurons promotes and inhibits the mouse feeding behavior, respectively^29, 30^. While previous studies have traced the axons of these two neurons at the population level^18, 19^, how individual *Agrp* or *Pomc* neurons innervate their downstream targets has yet to be examined by anterograde tracing of axons.

**Fig.5.**
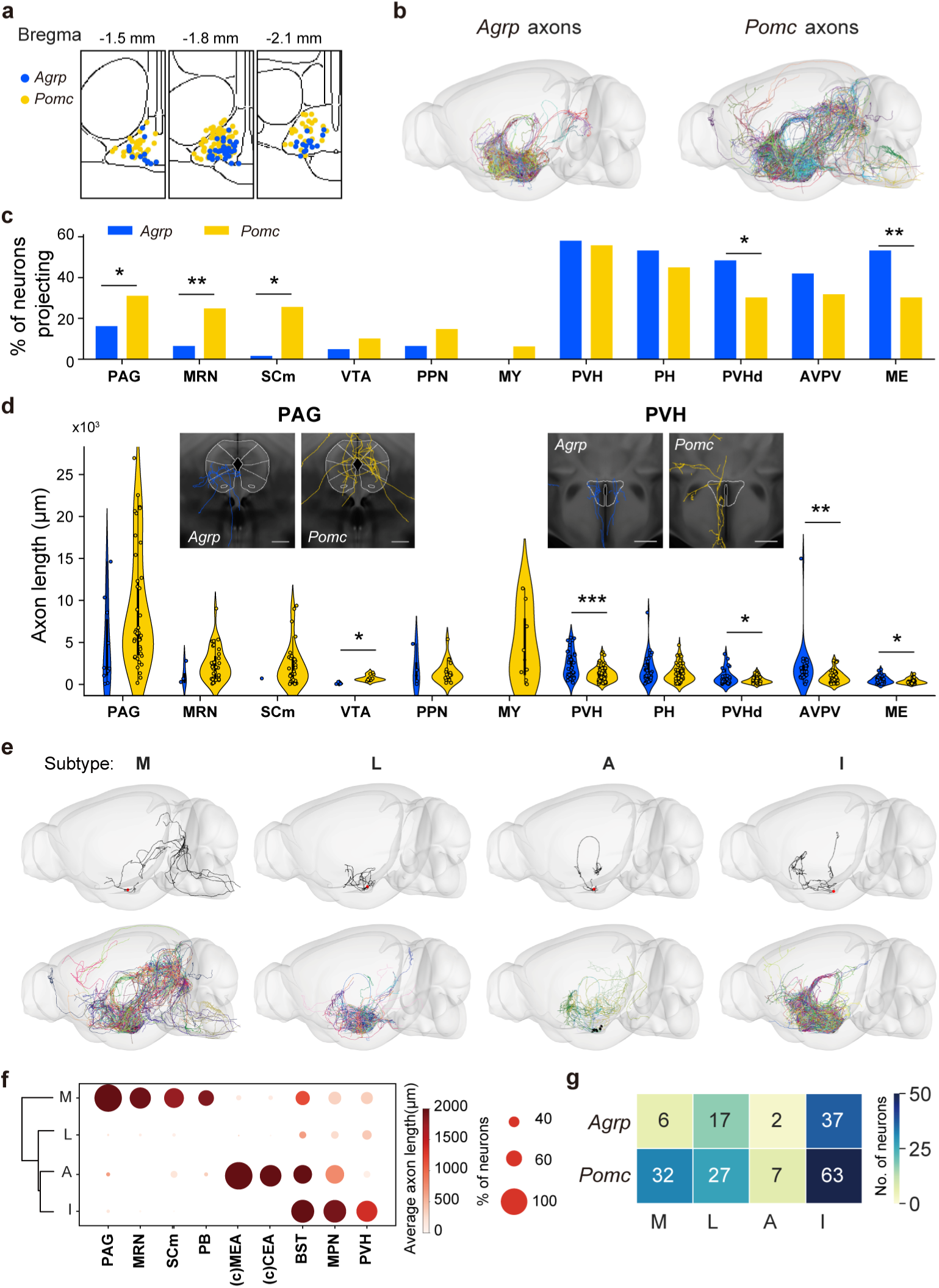
Arcuate *Agrp* and *Pomc* neurons differ in projectome subtypes. **a.** The soma location of the 62 *Agrp* neurons (blue dots) and 129 *Pomc* neurons (yellow dots) in the arcuate nucleus. **b.** The total axon projections of all *Agrp* (left) and *Pomc* neurons (right) throughout the brain. Each neuron was labeled with a different color. **c-d.** The percentage of neurons projected to (c) and the axon projection length of individual neurons (d) in indicated brain regions. Images in d show axon projections from a representative *Agrp* or *Pomc* neuron in PAG or PVH. Scale bar, 500 μm. **e.** Morphology-based clustering analysis revealed four projectome-defined subtypes of *Agrp* and *Pomc* neurons, termed “M”, “L”, “A” and “I”. The top panels show the projection of a representative neuron in each subtype, and the bottom panels show all axon projections of neurons in each subtype. **f.** A dot plot depicting selective projections of each subtype (row) in each brain area (column). The dot circle’s size and color intensity indicate the percentage of neurons in each subtype that projected to the indicated brain area and the average projection length (in μm) in a heatmap fashion, respectively, with the scale on the right. (c)MEA and (c)CEA indicate contralateral projection. **g.** The number of *Agrp* and *Pomc* neurons in each projection subtype. *See also Extended Data Fig.8*

Initial qualitative inspection of axon projections of all arcuate *Agrp* vs. *Pomc* neurons showed prominent projections of *Pomc* but not *Agrp* neurons to the midbrain, pons, and medulla regions (**Fig.5b**). This was further quantified by higher percentages and a stronger projection strength of individual *Pomc* neurons in PAG, MRN, SCm, and VTA (**Fig.5c-d**). By contrast, individual *Agrp* neurons showed more robust projections to hypothalamic targets, including PVH, PVHd, AVPV, and ME (**Fig.5d**). Moreover, morphology-based clustering of these *Agrp* and *Pomc* neurons yielded four subtypes, with subtypes “M”, “L”, “A”, and “I” characterized by strong projection to <u>M</u>idbrain areas, <u>L</u>ocally within the arcuate nucleus, medial and central <u>A</u>mygdala on the contralateral side, and Intra-hypothalamically, respectively (**Fig.5e-f**). Consistently, we also found a significant enrichment (*p = 0.035*) of “M” neurons in the *Pomc* population (**Fig.5g**).

In addition, we observed that individual *Agrp* neurons often innervated multiple targets through axon collaterals in various brain regions, including BST, LHA, PVH, PAG, PVT, CEA, and PB (Extended Data Fig.8). Our findings of extensive *Agrp* axon collateralization contrasted with previous retrograde tracing studies, which suggested that distinct and dedicated pools of *Agrp* neurons project to separate downstream targets with little co-innervation^31^. This discrepancy could be potentially attributed to the infection bias or the dependence on innervation strength in retrograde labeling methods. Taken together, our single-neuron projectome analysis revealed complex but divergent projection patterns of individual *Agrp* and *Pomc* neurons.

### Comparison of single-cell projectomes across hypothalamic subnuclei

Having established that individual peptidergic populations comprised multiple projectome-defined subtypes, we further examined whether regional differences in projectome-defined subtypes could shed light on functional differences of distinct hypothalamic areas, taking advantage of the full hypothalamus coverage of our dataset. Previous studies have shown that activating the medial preoptic area (mPOA) elicits male mating and parental care^32–35^, activating the ventrolateral division of VMH (VHMvl) promotes territorial and maternal aggression^36–40^, and activating the dorsomedial division of VMH (VMHdm) promotes predatory defense^41^ (**Fig.6a**). We postulated that differences in long-range axon projection patterns of individual neurons in these three regions might account for their functional differences in driving distinct behaviors.

**Fig. 6.**
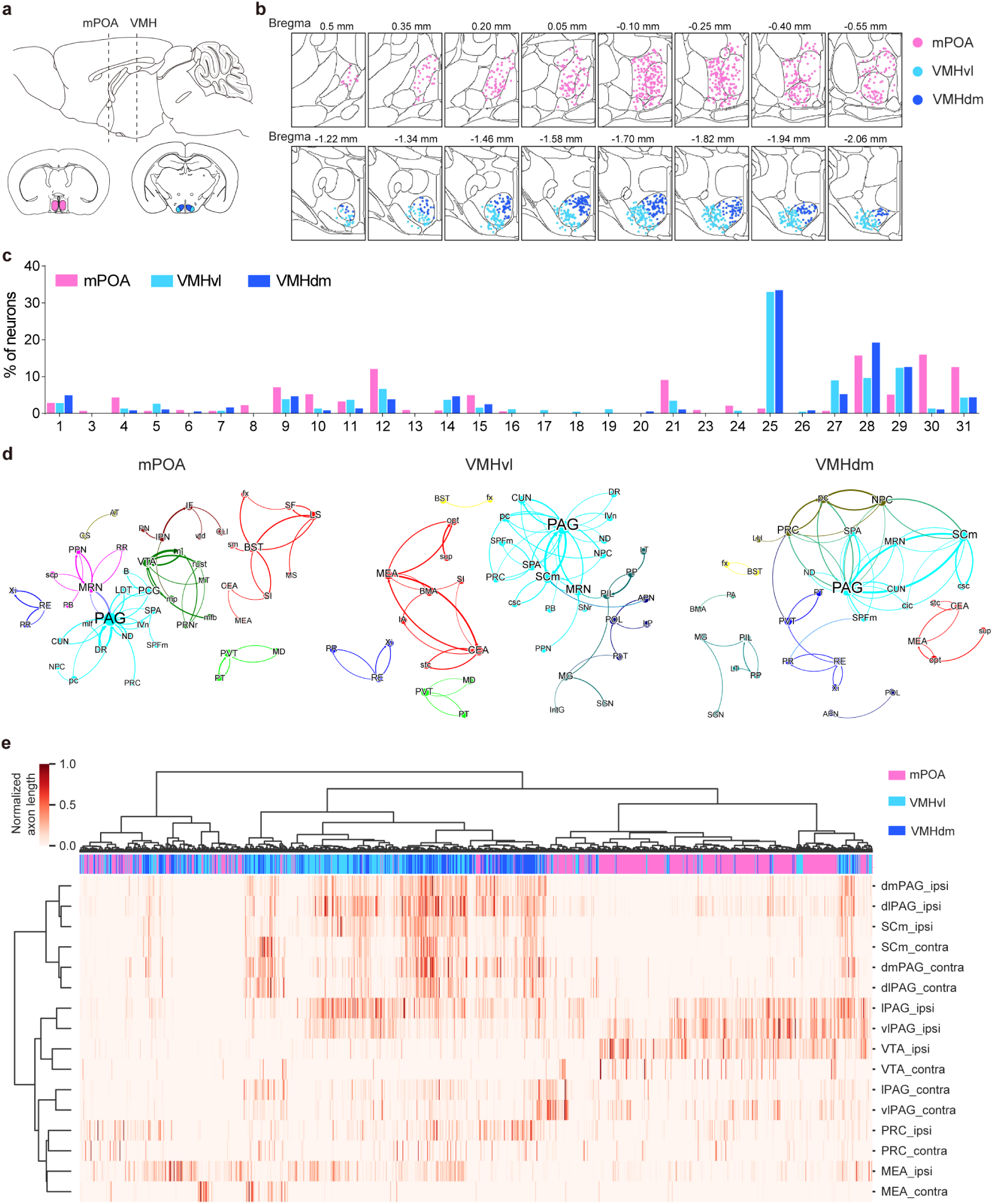
Comparison of single-cell projectome across hypothalamic subnuclei. **a.** Schematics showing the anatomic location of mPOA, VMHvl, and VMHdm in a sagittal (top) and coronal view of the mouse brain. **b.** The soma location of the 799 mPOA neurons, 470 VMHvl neurons, and 364 VMHdm neurons. **c.** The percentage of mPOA, VMHvl, and VMHdm neurons in the 31 projectome-defined subtypes. No neurons from these three regions were found in subtype #2 or #22. **d.** Graph theory-based network analysis of the extra-hypothalamic targets of individual mPOA, VMHvl, and VMHdm neurons. Colored nodes denote brain regions in the same module. **e.** A summary of normalized axon projection length in the ipsilateral (ipsi) and contralateral (contra) side of each brain area indicated on the right for each row. Each tick represents the normalized value of the axon projection length of a single neuron in a heatmap fashion with the scale shown upper left. Color strips above the heatmap represent each neuron’s location, with mPOA/VMHvl/VMHdm represented by pink/light blue/blue, respectively.

To this end, we manually demarcated the mPOA, VMHvl, and VMHdm areas within the Allen CCFv3 template and identified 799, 470, and 364 neurons in our dataset whose somata were localized in these three regions, respectively (**Fig.6b**). Interestingly, there was a strong regional bias in the distribution of projectome-defined subtypes, especially some midbrain-only projecting subtypes, across these three regions (**Fig.6c**). For example, subtypes #25 and #27, characterized by bilateral and contralateral co-innervation of PAG, preferred VMHvl and VMHdm; on the other hand, subtypes #30 and #31, with co-innervation of ipsilateral vlPAG and lPAG, favored mPOA (**Fig.6c**, **Fig. 3b-e**).

Next, we performed graph theory-based network analysis of the extra-hypothalamic targets of individual mPOA, VMHvl, and VMHdm neurons to pinpoint downstream targets critical for driving behaviors controlled by these hypothalamic neurons. For this analysis, we considered each target region as a node in the network and the axon trajectories passing through the target regions as directed edges between the nodes. Although neurons in the mPOA, VMHvl, and VMHdm regions had many similar downstream targets, as reflected by shared network nodes, there were remarkable differences in the network configuration, as shown by differences in the edges between nodes (**Fig.6d**). Most notably, a VTA node was present in the projection network of mPOA neurons but absent in those of VMHvl and VMHdm neurons; by contrast, SCm was a prominent node for VMHvl and VMHdm but not the mPOA network. Moreover, although VMHvl and VMHdm networks were more similar to each other than the mPOA network, they differed in several aspects. For example, the module comprising MEA and CEA nodes was more prominent in the VMHvl than in the VMHdm network.

These distinctions among mPOA, VMHvl, and VMHdm targeting networks were further corroborated by single-cell projectome analysis of individual mPOA, VMHvl, and VMHdm neurons (**Fig.6e**). We observed that mPOA neurons send strong ipsilateral projections to lPAG and vlPAG with frequent collateral projections to VTA. By contrast, VMHvl neurons tended to innervate all columns of the ipsilateral PAG with few collaterals to VTA. Meanwhile, VMHdm neurons showed strong bilateral projections to dmPAG, dlPAG, and SCm. These projection characteristics of individual neurons thus underlie the configuration features of the targeting network for each hypothalamic region. Furthermore, these results of targeting network analysis support the notion that VTA is a critical downstream target in promoting mating and parental care^34, 42^, whereas SCm and MEA are important for promoting predatory defense and territorial aggression, respectively^43–45^. We envision similar analysis could be performed for other hypothalamic neurons in the database to elucidate other neural circuits underlying diverse hypothalamic functions.

### Modular subnetwork organization of intra-hypothalamic projections

An additional outcome of our single-neuron projectome analysis was the elucidation of extensive intra-hypothalamic connectivity (see **Fig. 1f**), which is well-known but difficult to characterize by bulk tracing methods due to the small size and the densely-packed organization of the hypothalamus. In our dataset, the registered intra-hypothalamic projections covered 83.6% of the ipsilateral and 25.7 % of the contralateral volume of the hypothalamus (at the 10-μm voxel resolution), respectively, providing a comprehensive high-resolution view of the intra-hypothalamic connectivity. To further analyze intra-hypothalamic connectivity patterns, we divided the hypothalamus into 7425 cubes (100 μm * 100 μm * 100 μm), among which 3445 contained at least one soma. Using the Louvain algorithm previously described^23^, we constructed a network of intra-hypothalamic connectivity based on reciprocal projections between the cubes that contained both soma and axon (**Fig.7a**). We found that intra-hypothalamic connectivity could be characterized by six subnetworks enriched with recurrent intra-subnetwork connections (**Fig.7a**).

**Fig. 7.**
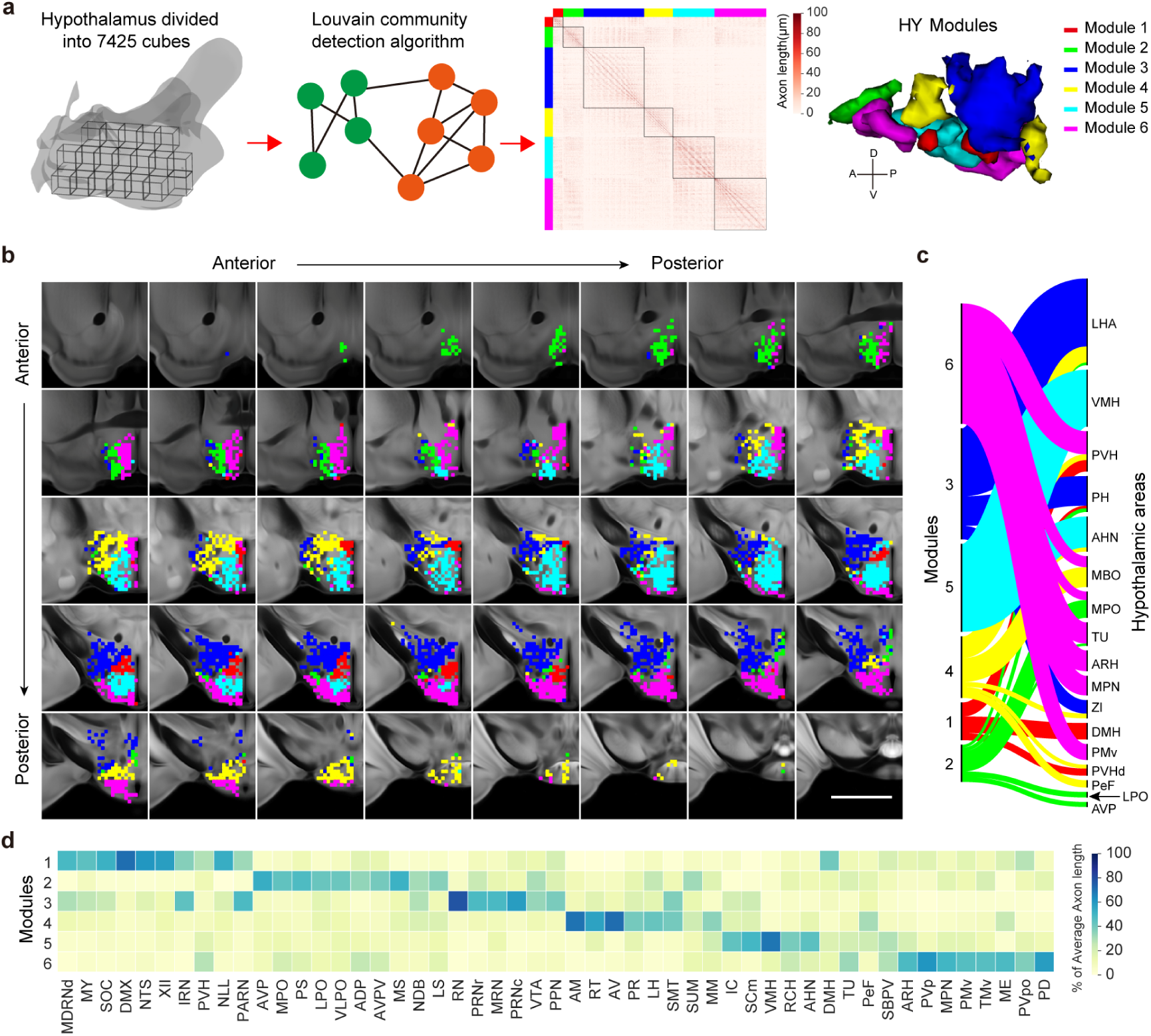
Modular subnetwork organization of intra-hypothalamic projections. **a.** Schematics illustrating the procedure to identify hypothalamic subnetwork modules enriched for intra-modular connectivity. Briefly, the brain volume of the hypothalamus was divided into 7425 cubes, and sets of cubes enriched for intra-cube reciprocal connections were identified through the Louvain community detection algorithm. In total, six hypothalamic subnetwork modules (color-coded) were identified. **b.** The six hypothalamic subnetwork modules are plotted in serial coronal sections spaced 100 μm apart. Scale bar, 500 μm. **c.** Correspondence between identified subnetwork modules and anatomically defined hypothalamic nuclei. **d.** Heatmap representing the percentage of the average axon length in a target area that were from each hypothalamic subnetwork module.

Mapping these subnetwork modules onto the brain template showed that each module corresponded to a distinct set of previously identified hypothalamic nuclei, with the prominence of a few: DMH and PVH in module 1, MPO and LPO in module 2, LHA and PH in module 3, LHA and MBO in module 4, VMH and AHN in module5, and MPN, ARH, and PMv in module 6 (**Fig.7b-c**). In support of the notion that these modular subnetworks may be functionally relevant, we found that modules 5 and 6 corresponded well to the known hypothalamic “defensive” and “reproductive” networks. Moreover, module 6 also placed ARH in the “reproductive” network of the hypothalamus **(Fig.7b-c)**. The functional significance of these modules was further supported by the distinct preference for the downstream targets displayed by each module (**Fig.7d**). We surmise that this modular subnetwork of intra-hypothalamic connections may organize hypothalamic outputs for regulating physiological functions and innate behaviors.

## Discussion

We have built an extensive dataset of whole-brain projectomes of ∼7000 peptidergic neurons from nearly all hypothalamic regions and identified 31 projectome-based subtypes. A critical point demonstrated by the present study is the lack of one-to-one correspondence between projectome-defined and transcriptome-defined subtypes. We found that the expression of neuropeptides, which sometimes mark individual transcriptome-defined excitatory or inhibitory neuron types, was always found in multiple projectome-defined subtypes. Conversely, each projectome subtype expressed multiple neuropeptides of distinct combinations with preferential enrichment of a few. Similar combinatorial correspondence between projectome- and transcriptome-defined subtypes was also found for PFC projection neurons^23^. Thus, axon projection and gene expression patterns reflect different facets of neuronal features that are likely to be interdependent. Moreover, by analyzing single-neuron projectomes for multiple molecular markers across distinct hypothalamic regions, we may gain insights into neural circuit mechanisms underlying the diverse and specific hypothalamic functions mediated by different neuronal types.

Our analyses revealed extensive axon collateralization and arborization in some projectome subtypes that enable a single hypothalamic neuron to simultaneously innervate multiple brain areas, corresponding areas in both hemispheres, and distinct subdomains of a given area. Previous studies using conventional retrograde tracer or viral tracing strategy have identified hypothalamic neurons that appeared to project to single prominent downstream targets^31, 32^. However, our anterograde tracing method that directly labels the axon arbors throughout the whole brain projection rarely revealed single-target axon projection. Noteworthily, our analysis used total arbor length or density instead of synaptic site counts over an area to define the innervation strength at the target. This is because the synaptic counts tended to be directly proportional to the arbor length (Supplementary Table 4). Furthermore, neuropeptides are likely released from non-synaptic sites along the axon arbor^46^. Moreover, our conclusion of extensive axon collateralization was confirmed by the analysis of hypothalamic neurons previously reported by the MouseLight project^25^ (Extended Data Fig.4). Thus, extensive axon collateralization likely represents a general principle of hypothalamic projections, which may facilitate coordinated downstream actions of hypothalamic projection neurons. Whether or how co-release of classical neurotransmitters with hypothalamic neuropeptides, such as glutamate for *Orexin* and *Pomc* neurons or GABA for *Agrp* neurons, in selected target subdomains could regulate various brain functions remains to be studied.

Additionally, while extensive intra-hypothalamic connectivity has been shown previously^47^, our study provided important new information in characterizing the organization patterns of intra-hypothalamic connectivity. We found six modular subnetworks enriched with intra-modular reciprocal projections, implicating organized cross-talks among hypothalamic peptidergic neurons. Two of the six identified modules roughly matched the previously known hypothalamic “defensive” and “reproductive” networks^48–50^, supporting the functional relevance of these modular subnetworks. Whether neurons in the modules are synaptically connected remains to be examined. Modular subnetworks generally allow localized information processing before more global integration^51^. Identifying modular subnetworks within the hypothalamus indicates potential functional specializations of distinct hypothalamic subdomains, which may facilitate the instantiation of internal brain states^52^, such as arousal or emotion states, associated with hypothalamic activation.

Finally, considering the hypothalamus as an example of subcortical structures, the present work illustrates the complexity of both long-range projections and local connections and their potential relevance for understanding neural circuit functions of the subcortical structure. As our single-neuron tracing is restricted to axon projections, mapping the input connections to each hypothalamic neuron subtype is required to delineate the complete connectivity of the neuron within the neural network, which is necessary to understand their functional roles fully in the future. Further integration of transcriptomic and other x-omic information into the connectivity-defined subtypes will allow complete characterization of distinct neuronal subtypes marked by molecular and connectivity features. To accomplish this goal, we advocate using a consensual annotation framework^24^ to facilitate the integration of information generated in different labs. These analyses of neuronal subtypes, coupled with immediate early gene analysis and functional manipulation, would eventually reveal how coordinated hypothalamic neuronal activities give rise to behavioral and physiological changes in the body.

## Methods

### Animals

Genetically modified mouse lines including *Adcyap1-2A-Cre* (B6.Cg-Adcyap1^tm1.1(cre)Hze^/ZakJ, Cat#030155), *Agrp-IRES-Cre* (STOCK Agrp^tm1(cre)Lowl^/J, Cat# 012899), *Avp-IRES-Cre* (B6.Cg-Avp^tm1.1(cre)Hze^/J, Cat# 023530), *Crh-IRES-Cre* (B6(Cg)-Crh^tm1(cre)Zjh^/J, Cat#012704), *Nts-IRES-Cre* (B6;129-Nts^tm1(cre)Mgmj^/J, Cat# 017525), *Oxytocin-IRES-Cre* (B6;129S-Oxt^tm1.1(cre)Dolsn^/J, Cat# 024234), *Pdyn-IRES-Cre* (B6;129S-Pdyn^tm1.1(cre)Mjkr^/LowlJ, Cat#027958), *Penk-IRES-Cre* (B6;129S-Penk^tm2(cre)Hze^/J, Cat# 025112), *Pmch-Cre* (STOCK Tg(Pmch-cre)1Lowl/J, Cat# 014099), *Pomc-Cre* (STOCK Tg(Pomc1-cre)16Lowl/J, Cat# 005965), *Sst-IRES-Cre* (STOCK Sst^tm2.1(cre)Zjh^/J,Cat# 018973), *Tac1-IRES-Cre* (B6;129S-Tac1^tm1.1(cre)Hze^/J, Cat# 021877), *Tac2-Cre* (B6.129-Tac2^tm1.1(cre)Qima^/J,Cat# 018938), *Trh-IRES-Cre* (B6;129S-Trh^tm1.1(cre)Mjkr^/LowlJ, Cat# 032468), *Vip-IRES-Cre* (STOCK Vip^tm1(cre)Zjh^/J, Cat# 010908) were purchased from the Jackson Laboratory and bred in house. Wildtype mice of the C57BL/6J background were purchased from Slac Laboratory. All mice were housed on a 12 h light / dark cycle with water and food *ad libitum* in the institute’s animal facility. Only adult males over 8 weeks old were used in the study. All experimental protocols were approved by the Animal Care and Use Committee of the Center for Excellence in Brain Science and Intelligence Technology, Chinese Academy of Sciences, Shanghai, China (IACUC No.NA-016-2016)

### Virus

We used various methods to achieve sparse labeling neurons with cell-type specificity. For virus injected into *Cre* animals, these methods included diluting AAVs expressing *Cre*-dependent green fluorescent proteins of different versions, or mixing AAVs expressing *Cre*-dependent *FlpO* and AAVs expressing *Flpo* dependent fluorescent protein at a ratio. Viral mixing was achieved by either combining two AAVs produced separately or by co-transfection of mixed plasmids for both AAVs to cells during the viral production step as described ^53^. In addition, to label *Orexin* neurons, we injected AAVs encoding mNeoGreen under the control of the *Orexin* promoter (PPORX) into wildtype males.

The following virus were used in this study: AAV-CAG-DIO-FlpO (titer, 5.0*10^9^ genomic copies / mL) were purchased from Shanghai Taitool Bioscience, Co. Ltd. AAV-EF1a-fDIO-EYFP (titer, 1.31*10^13^ genomic copies / mL); AAV-EF1a-DIO-Ypet-2A-mGFP (titer, 2.82*10^12^ genomic copies / mL); AAV-EF1a-DIO-FlpO (titer, 3.2*10^9^ genomic copies / mL or 4.1*10^9^ genomic copies / mL or 1*10^9^ genomic copies / mL or 4.0*10^10^ genomic copies / mL,1.25*10^9^ genomic copies / mL); AAV-EF1a-fDIO-Ypet-P2A-mGFP (titer, 7.4*10^12^ genomic copies / mL or 6.3*10^12^ genomic copies / mL or 9.8*10^12^ genomic copies / mL or 2.31*10^12^ genomic copies / mL or 1.05*10^12^ genomic copies / mL or 5.5*10^12^ genomic copies / mL). The JS series of virus were a mixture of AAV-hSyn Con/Fon EYFP and AAV-EF1a-FlpO generated at different ratio during the virus package step (1:4000, titer, 5.2*10^12^ genomic copies / mL; 1:8000, titer, 1.41*10^13^ genomic copies / mL; 1:40000, titer, 5.1*10^12^ genomic copies / mL); AAV-PPORX-mNeoGreen (titer, 1.0*10^13^ genomic copies / mL); AAV-EF1a-DIO-EYFP (titer, 1.0*10^12^ genomic copies / mL) were produced by the Gene Editing Core Facility at the institute.

### Surgery

Stereotaxic surgeries were performed to inject AAVs to the subregion of the hypothalamus according to the Paxinos and Franklin Mouse Brain Atlas (2nd edition) as previously described. The detailed information of AAVs injected, injection coordinates, and injection volume for each brain sample was listed in Supplementary Table 1.

#### Histology

Animals were anesthetized with 1% pentobarbital sodium (50 mg/kg, i.p.; Merck) and perfused with DEPC-treated PBS followed by ice-cold 4% PFA in PBS. Brains were post-fixed in PFA overnight at 4 °C and dehydrated with 30% sucrose in DEPC-PBS. Afterward, brains were sectioned at 20 μm thickness and mounted onto SuperFrost Plus Slides (Fisher Scientific, Cat# 12-550-15). After drying in the air, brain sections were stored in −80 °C. The in situ hybridization was performed using the RNAscope kit (ACD Bio.), following the user manual. Probes used in the experiment against specific peptide mRNA were ordered from ACD Bio. After in situ process, brain sections were blocked in 2.5% BSA (Sigma Cat #V900933) for one hour and then stained overnight at 4 °C with primary antibody Chk pAb to GFP (Abcam Cat# ab13970, dilution 1:300). In the next day, the brain sections were rinsed three times with 1×PBS, then incubating with the secondary antibody, goat-anti-chicken Alexa Fluor 488 (Jackson Immuno Research Laboratories, Cat# 103-545-155, dilution 1:300) for 2 hours. After that, brain sections were rinsed three times with 1×PBS, and finally counter-stained with DAPI (Sigma, Cat# d9542, 5mg/ml, dilution 1:1000). Images were captured under a 20/40X objective using a confocal microscope (Nikon C2/Olympus FV3000).

### fMOST imaging, tracing and registration

#### Imaging

The fMOST imaging was acquired as previously described ^23^. Briefly, the dissected brains were post-fixed in 4% paraformaldehyde and then embedded in Lowicryl HM20 resin (Electron Microscopy Sciences, 14340). The resin-embedded brains were imaged in a water bath containing propidium iodide (PI) under an fMOST microscope at a voxel resolution of 0.32 µm × 0.32 µm × 1 µm. Briefly, the sample surface in a coronal plane was imaged for GFP (for neuron tracing) and PI channels (for brain registration), then removed at 1 µm step by a fixed diamond knife. The imaging-sectioning cycle was continued until the entire brain sample was fully imaged.

#### Tracing

A new software tool, Fast Neurite Tracer (FNT), was developed in-house to trace long-range axon projection in tera-bytes datasets generated by light microscopy ^23^. Briefly, the original image data are split into smaller 3D data cubes by a program called “slice2cube” in FNT. At any time, about eight neighboring cubes around a position are loaded into computer memory automatically and visualized in 3D for tracing. The tracing process involves finding a putative path, examining the path, and extending the current neurite tree. At each step, confirmation from a user was required to proceed, thereby ensuring the correctness of tracing. An axon projection can be traversed in this way until completion. Different neurons in the same sample were traced by different personnel in parallel and cross-validated after completion. In addition, each traced neuron was independently checked by another person to ensure accuracy and each brain sample was subjected to random post-tracing quality check.

#### Registration

The whole brain with all reconstructed neurons’ information was registered into the standard Allen CCFv3 ^24^ using a previously described method ^23^. Briefly, we segmented several brain regions as landmarks through cytoarchitecture references. We performed diffeomorphic transformation and symmetric image normalization in Advanced Normalization Tools (ANTs) to acquire transformation parameters based on these landmarks. These transformation parameters were applied to all the traced neurons within the brain sample to remap the reconstructed neuron onto the Allen brain template.

### Data analysis

#### Neuron exclusion

All neurons were manually checked to ensure that they were correctly traced. Neuron found to have erroneous tracing data were excluded from further analysis. Additionally, neurons whose soma location were outside the hypothalamus were also excluded. In total, we included 7180 neurons in our analysis. We mirrored all neurons to the same hemisphere before the analysis procedures.

#### Parameter calculation

Base on the standard Allen CCFv3 annotation file (http://download.alleninstitute.org/informatics-archive/currentrelease/mouse_ccf/annotation/ccf_2017/), we calculated the *soma location*, *axon length* and *terminal number* for each neuron in each brain region using a self-developed neuron visualization and analysis software – *NeuronView* (https://gitee.com/bigduduwxf/NeuronView). To show the data in a more reader-friendly way, we manually combine some small brain areas to bigger ones for plotting. To calculate other parameters including Max Distance from soma, Max EucDistance from soma, Mean Contraction of each edge, Mean Partition asymmetry and Mean Fractal Dim we used L-measure^54^ (http://cng.gmu.edu:8080/Lm/help/index.htm).

#### Neuron visualization

All 3D neuron showcase were plot using a self-developed python package – *neuron-vis* (https://gitee.com/bigduduwxf/neuron-vis). Colors were randomly assigned to individual neurons plotted at a population level. For zoom-in views of the axon projections in particular areas in a 2D view, we picked 3 neurons in each subtype that had the highest axon length in chosen brain region and plotted their axons on the center coronal section of the brain region in the Allen template brain.

#### Projectome-based neuron classification

We calculated a modified Hausdorff match distance^55^ as the similarity index of neuron pairs in our dataset. Briefly, we broke each neuron traces into sets of evenly spaced points in the 3D Allen template brain space to generate two sets of points for a neuron pair, A and B. For a given point “i” in the set A, we then calculated all the distances to every single point in the set B and picked out the smallest value, i.e. the closest distance to neuron B. This value is defined as d (Ai, B). We re-iterated this process for all points on neuron A, generating a collection of d (Ai_1→j,_ B), defined as D (A, B), from which we calculated the mean and the max value of the collection. We then calculated a weighted average by taking into the consideration of the ratio, α, of points in the set A that had d (Ai, B) smaller than the mean value. Thus d (A → B) was directional A → B distance and was calculated as α * Mean (D (A, B) + (1-α) * Max (D (A, B)). The directional distance B → A was calculated in the same manner. The similarity index between neuron pair A and B is the mean of d (A → B) and d (B → A). The formulas were as following:

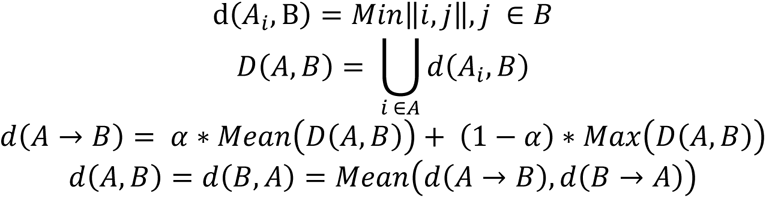

After generating the similarity index of all neuron pairs, we performed hierarchical clustering of the matrix using Ward’s linkage to identify projectome-defined neuron subtypes. The clustering threshold was at a distance equaling to or smaller than 8000. This yielded 31 projectome-defined subtypes.

#### Validation of image registration

To calculate the registration accuracy, we randomly picked four brain samples and manually labelled three brain areas in the 3D space, fi, ME and IPN, before the registration. We then got the coordinates of these brain areas after registration, calculated the mass center, and measured their Euclidean distance to the mass center of fi, ME and IPN as defined by the standard Allen brain template, to illustrate the registration accuracy.

#### Selection of selective target areas for each subtype

To select the brain areas that were preferentially targeted by a given projectome-defined subtype in dot plots, we compared the percentage of neurons in each subtype and the number of neurons in the remaining subtypes that project to a given brain area by Fisher’s exact test. We chose the target brain areas the showed most significant differences for each subtype to plot.

#### PAG contralateral/ipsilateral preference index

Briefly, we first calculated the total axon projection lengths of the ipsilateral and contralateral PAG, and then calculated the contralateral/ipsilateral preference index using the following formula:

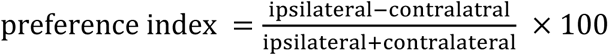

Neurons with the preference index >30, or <-30, or in-between were defined as ipsilaterally, contralaterally, and bi-laterally projecting, respectively.

#### PAG column/nucleus preference

Only axon projections in the ipsilateral PAG were considered. We first calculated axon projection density in each PAG column/nucleus and in the whole PAG by dividing the total axon projection lengths in each region by the corresponding volume. We then divided the axon projection density for each column/nucleus by that for the whole PAG to get to preference score for each domain.

#### PAG subdomains defined by hypothalamic co-innervation

In the standard Allen brain template, the PAG was not subdivided to columns as in the Paxinos and Franklin Mouse Brain Atlas. We therefore manually annotated the PAG into four columns, dmPAG, dlPAG, vlPAG and lPAG, in the standard Allen brain template using the Paxinos and Franklin Mouse Brain Atlas as the reference. We labeled the anterior section of the PAG that was not included in the four columns as the aPAG. To analyze co-innervation patterns of individual hypothalamic neurons, we first divided the ipsilateral PAG into 2117 cubes of 100 µm × 100 µm × 100µm size. For each PAG-projecting neuron, we calculated their axonal project length in each of 2117 these cubes. After obtaining the projection strength matrix, we conducted hierarchical clustering using Spearman’s rank correlation coefficient as the distance measure between neuron pairs and Ward’s linkage to obtain clustered cubes receiving highly correlated hypothalamus inputs. This method identified 7 PAG subdomains.

### Graph-theory-based analysis of target network

We constructed the adjacency matrix for directed weighted graphs using the routes of each neuron to determine the community characteristics of the target regions of mPOA, VMHvl, and VMHdm neurons. We traversed each neuron’s points from soma to the terminal and extracted the projection source region A for each target region B, labeling the connection as A->B = 1. With each neuron having its own directed adjacency matrix, we summed these matrices for mPOA, VMHvl, and VMHdm neurons to obtain the final adjacency matrix for each region. The target areas served as nodes, with area-area connections as edges, downstream connections as direction, and the number of neurons with such connections as edge weight. To eliminate outliers, we only kept nodes with a projection density greater than 20% of the population’s non-zero projection density (density>= 4.29 µm/ mm^3^) and connections shared by more than 20% of the neurons. We also excluded the hypothalamus regions and regions with connections to less than 20% of all nodes in the network from our analysis for the simplicity of our network. We then performed modularity-based community detection ^56^ to better determine the internal structures of these target regions.

#### Modular subnetwork organization of intra-hypothalamic projections

To analyze the intra-hypothalamic projection patterns, we divided the hypothalamus volume into 7425 cubes of 100 µm × 100 µm × 100 µm size considering only the hemisphere where the soma of all neurons was located. Of these 7425 cubes, 3445 had both neuron and axon projections in it. For each neuron, we calculated their axonal length in these 3445 cubes. To investigate the modular structure of connectivity, we used the Louvain community detection algorithm from the Brain Connectivity Toolbox ^57^ (https://sites.google.com/site/bctnet/) to find a consistent modular structures within these cubes. The final modular structure was obtained at γ = 1, with 6 intra-hypothalamus modules identified.

## Statistical analysis

Statistical analysis was performed in GraphPad Prism 7 or Python. Two-sided unpaired t-test and Fisher’s exact test were used to test for statistical significance. Statistical parameters including the exact value of n and statistical significance are reported in the text and in the figure legends. The significance threshold was 0.05 (*, P < 0.05; **, P < 0.01; ***, P < 0.001).

## Acknowledgements

This work was supported by the Shanghai Municipal Science and Technology Major Project (grant No. 2018SHZDZX05), the Lingang Laboratory (grant No. LG202104-01-01, No. LG202104-01-04), National Science and Technology Innovation 2030 Major Program (grant No. 2021ZD0204400, 2021ZD0203200-03, 2021ZD0201000).

## Author contributions

Sample preparation: X. Ding, Z. Yu, M. Li, M. Hao, H. Zhou, X. Cao, S. Li, C. Wang, E Li, Y. Hu, Z. Tao, H. Li, X. Yu, M. Xu, H. Chang, Y. Zhang, H. Xu; fMOST imaging, image pre-processing and quality control: H. Gong, Q. Luo, A. Li, T. Jiang, J. Qi, X. Jia; Data processing and quality control: Z. Feng, B. Ren, Y. Chen, X. Shi, D. Wang, X. Wang, L. Han, Y. Liang, X. Wang; Data Analysis: Z. Jiao, T. Gao, W. Zhang, N. Biglari, E. E. Boxer, L. Steuernagel, S. M. Sternson, J. C. Brunning, M. Poo, D. J. Anderson, Y. Sun; Manuscript Writing: R. Stoop, M. Poo, X. Xu.

## Data and Code availability

All relevant data and code for this study can be made available by the Lead Contact upon reasonable request.

## Materials Availability

All unique/stable reagents generated in this study are available upon request.

## Extended Data

**Extended Data Fig. 1.**
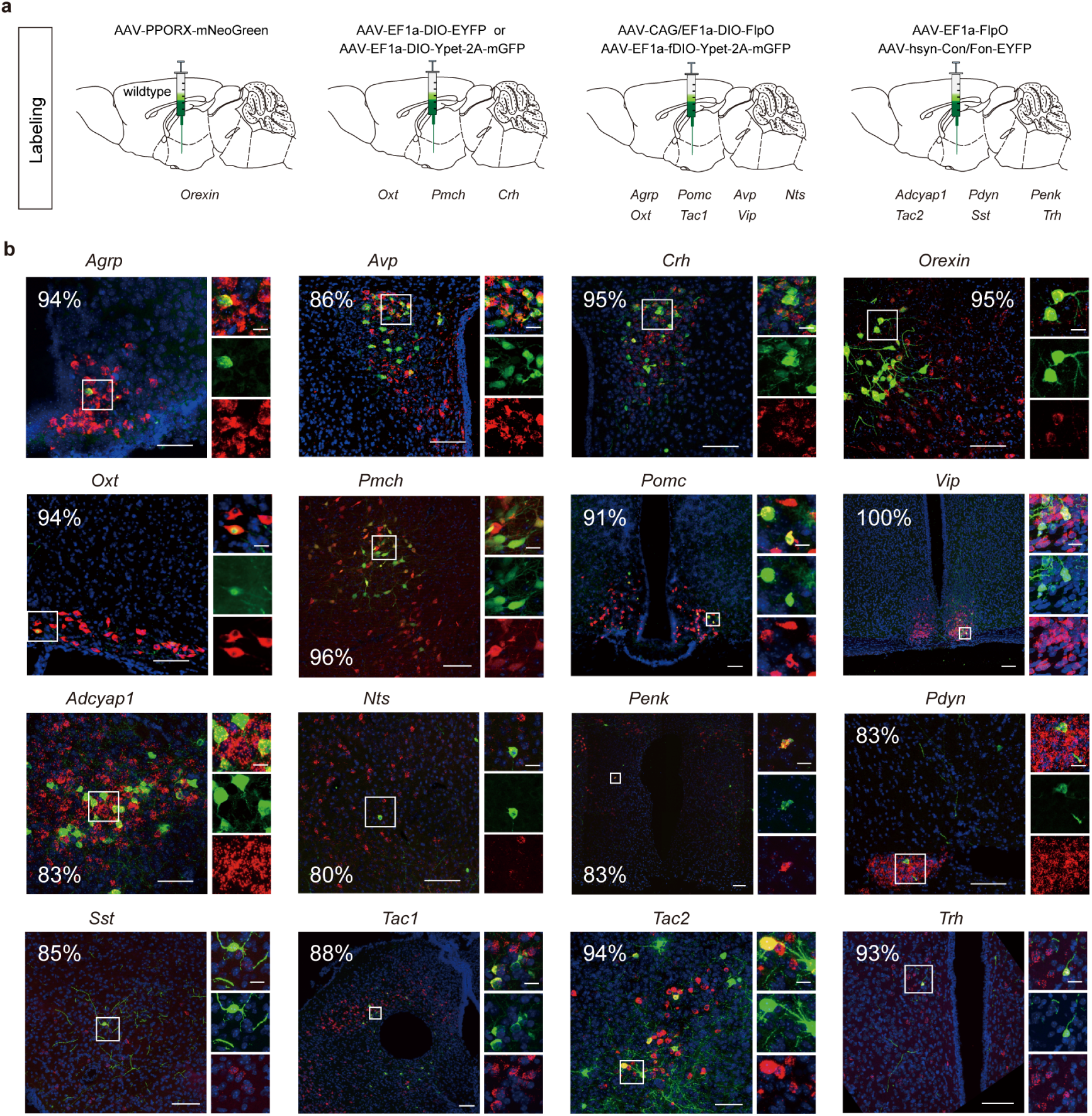
Viral strategies for sparsely labeling hypothalamic peptidergic neurons, *related to Fig.1*. **a.** To sparsely label neuropeptide-expressing populations listed at the bottom, the virus or virus mixture listed was injected into wildtype mice (for *Orexin*-expressing neurons) or *Cre*-expressing mice for all other peptidergic populations. **b.** Representative images showing co-localization of EYFP or GFP signal with *in situ* or immunohistochemistry signals of the indicated neuropeptide (red). The quantification shows the estimated percentage (%) of EYFP or GFP labeled cells that co-expressed the neuropeptide. The images on the right highlight regions within the white box on the left. Scale bar, left, 100 μm; right, 20 μm.

**Extended Data Fig. 2.**
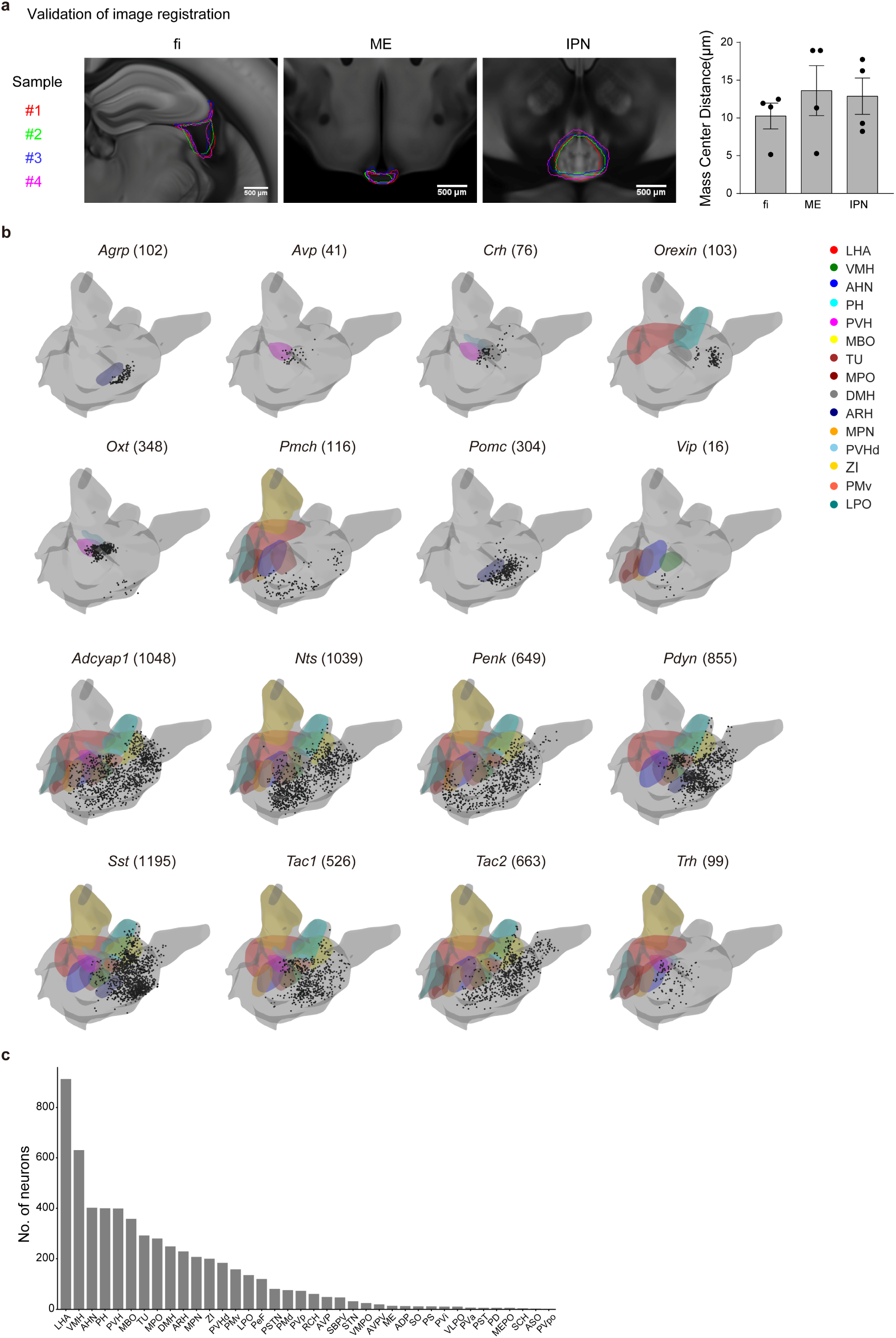
Brain registration and soma distribution of each peptidergic population, *related to Fig. 1*. **a.** Validation of image registration across different brain samples. Left, representative images showing the dispersion of the indicated structure (*fi*, ME, IPN) from four brain samples on the registered brains. Right, quantification of mass center distance (see Method). Data are presented as mean ± SEM. **b.** 3D view of the soma distribution of each of the 16 peptidergic neurons in various hypothalamic nuclei (outlined on the contralateral side, color-coded). The number of reconstructed neurons for each peptidergic population was indicated within the parenthesis. **c.** A bar graph depicting the number of reconstructed neurons with soma in the indicated hypothalamic nuclei.

**Extended Data Fig. 3.**
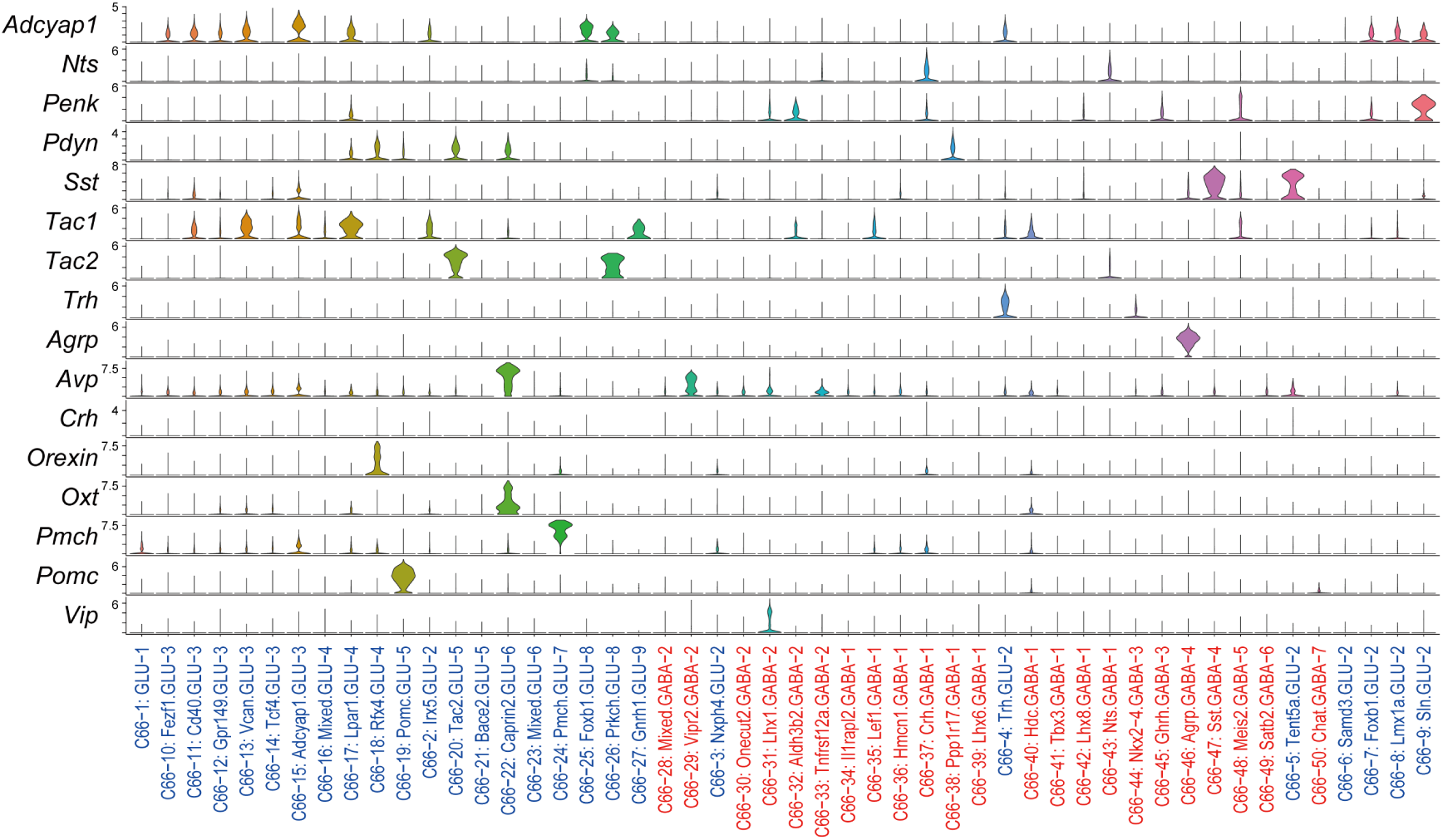
Distribution of neuropeptide expression in 66 neuronal types defined by scRNA-seq in the mouse hypothalamus, *related to Fig. 1*. Each column indicates a scRNA-seq-defined neuron type, with blue labeling indicating excitatory and red inhibitory neurons. *Orexin, Oxt, Agrp, Pmch, Pomc, and Vip* appear to label a single transcriptionally defined neuronal type.

**Extended Data Fig. 4.**
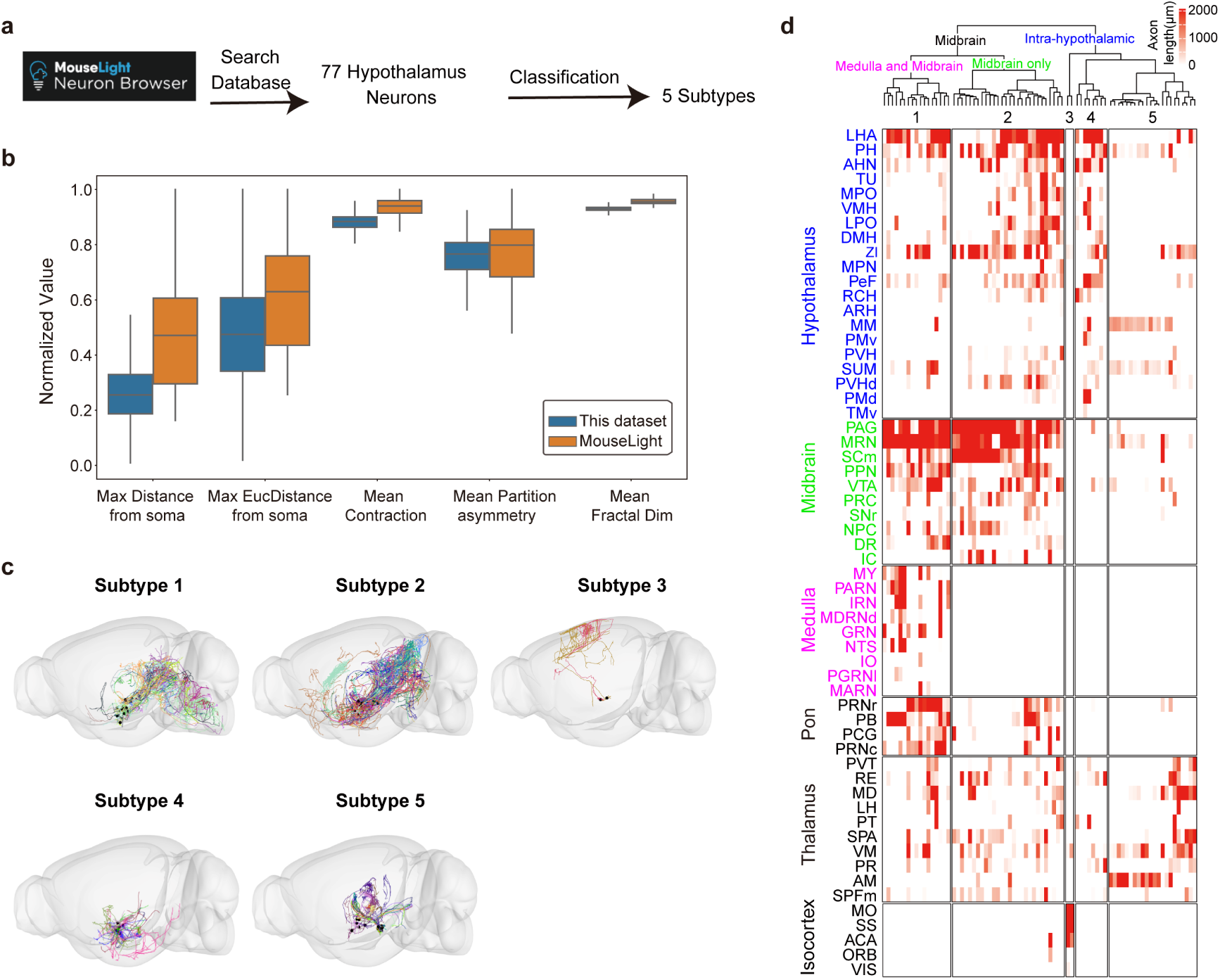
Morphology-based clustering analysis of 77 hypothalamic neurons reported previously, *related to Fig. 1.* **a.** We searched the published dataset and found 77 neurons whose soma location was within the hypothalamus. We classified these 77 neurons using the same algorithm described in Fig. 1. **b.** Several morphological parameters appear comparable between the 77 and 7180 neurons we reconstructed. Data are presented as median (center line) with 25/75 percentile (box) and min and max of the data points (whiskers) **c.** We clustered these 77 neurons into five projectome-defined subtypes. The total axon projections of neurons in each subtype were plotted in a 3D brain. **d.** A summary of axon projection length (in μm) of the given projectome-defined subtypes, divided into three major classes, in each brain area (color-coded, left). Each tick represents the axon projection length of a single neuron in each area in a heatmap fashion with the scale shown above.

**Extended Data Fig. 5.**
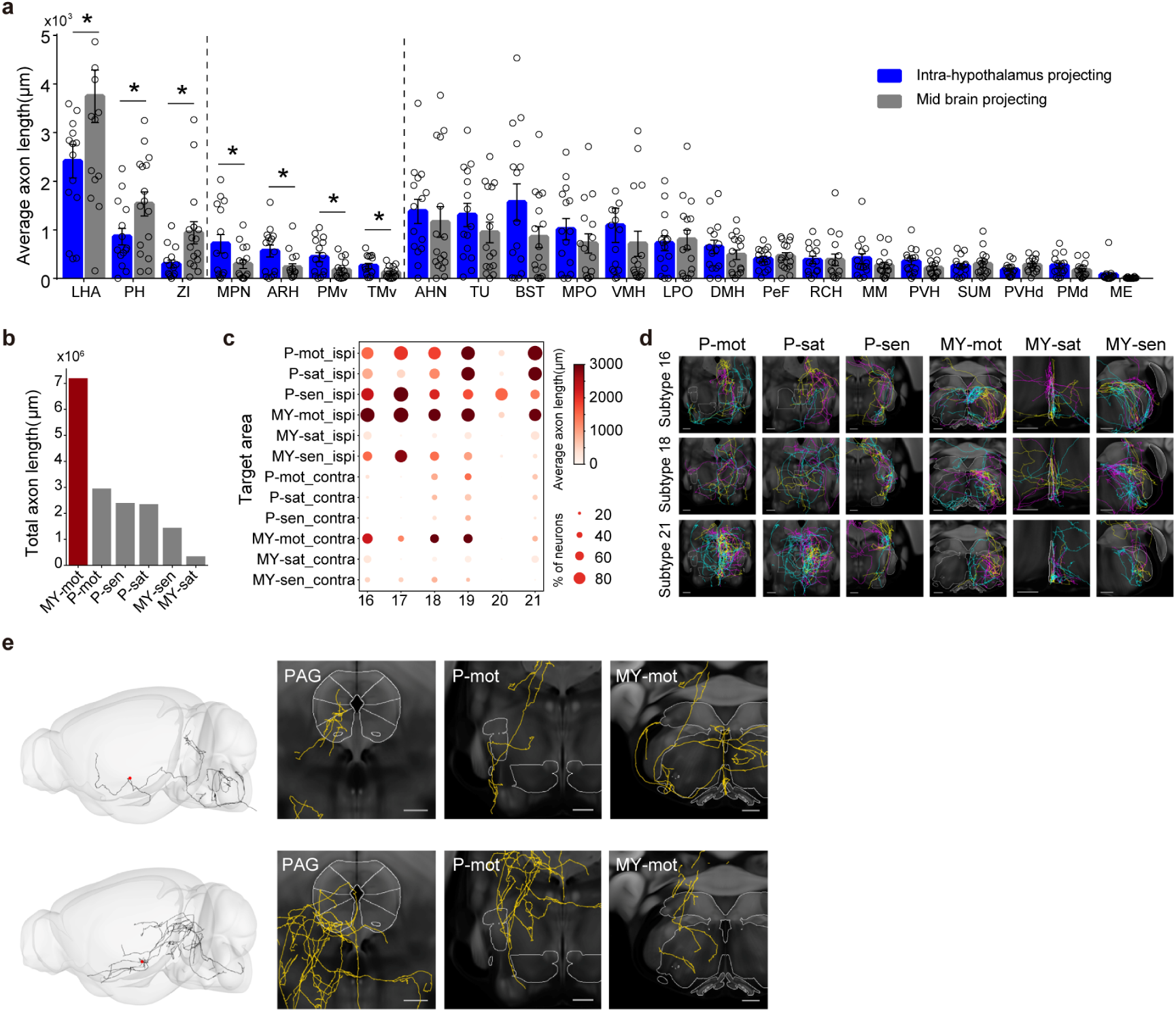
Characteristics of midbrain-projecting hypothalamic peptidergic neurons, particularly the medulla and midbrain class, *related to Fig. 3*. **a.** Comparison of the average axon projection length in the indicated hypothalamic nuclei for neurons in the intra-hypothalamic projecting subtypes (#1 - #15) versus those in the midbrain-projecting subtypes (#16 - #31). Data are presented as mean ± SEM. Each circle represents one subtype. **b.** The total axon projection length of medulla and midbrain-projecting class (subtypes #16-#21) in various medulla and pon areas. **c.** A dot plot of the percentage of neurons within each subtype (column) projecting to the indicated areas in pons or medulla on the ipsilateral or contralateral side (row). The color intensity within the circle indicates the average projection length emanating from neurons of each subtype. Scales are shown on the right. **d.** Representative images showing axon projection of neurons of the indicated subtypes (left) in the labeled brain areas (top). Scale bar, 500 μm. **e.** A 3D view of axon projections of two representative medulla and midbrain-projecting neurons on the left with images on the right showing axon collaterals in indicated brain areas.

**Extended Data Fig. 6.**
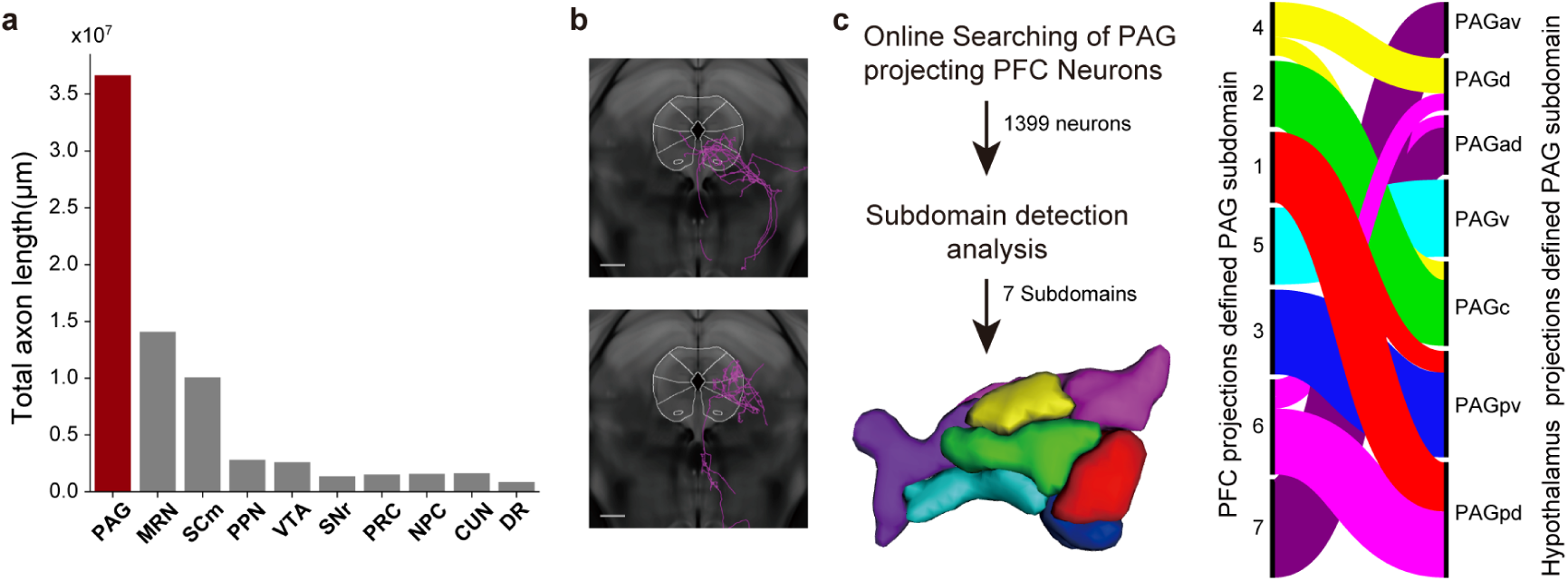
PAG subdomains defined through analysis of single-neuron projectome, *related to Fig. 3.* **a.** Bar graphs showing the total axon projection length of midbrain-projecting neurons (subtype #16 - #31) in various midbrain areas. **b.** Representative images showing strong PAG column/nucleus preference of projecting neurons. **c.** Correspondence between PAG subdomains defined by analysis of PFC or hypothalamus single-neuron projectomes.

**Extended Data Fig. 7.**
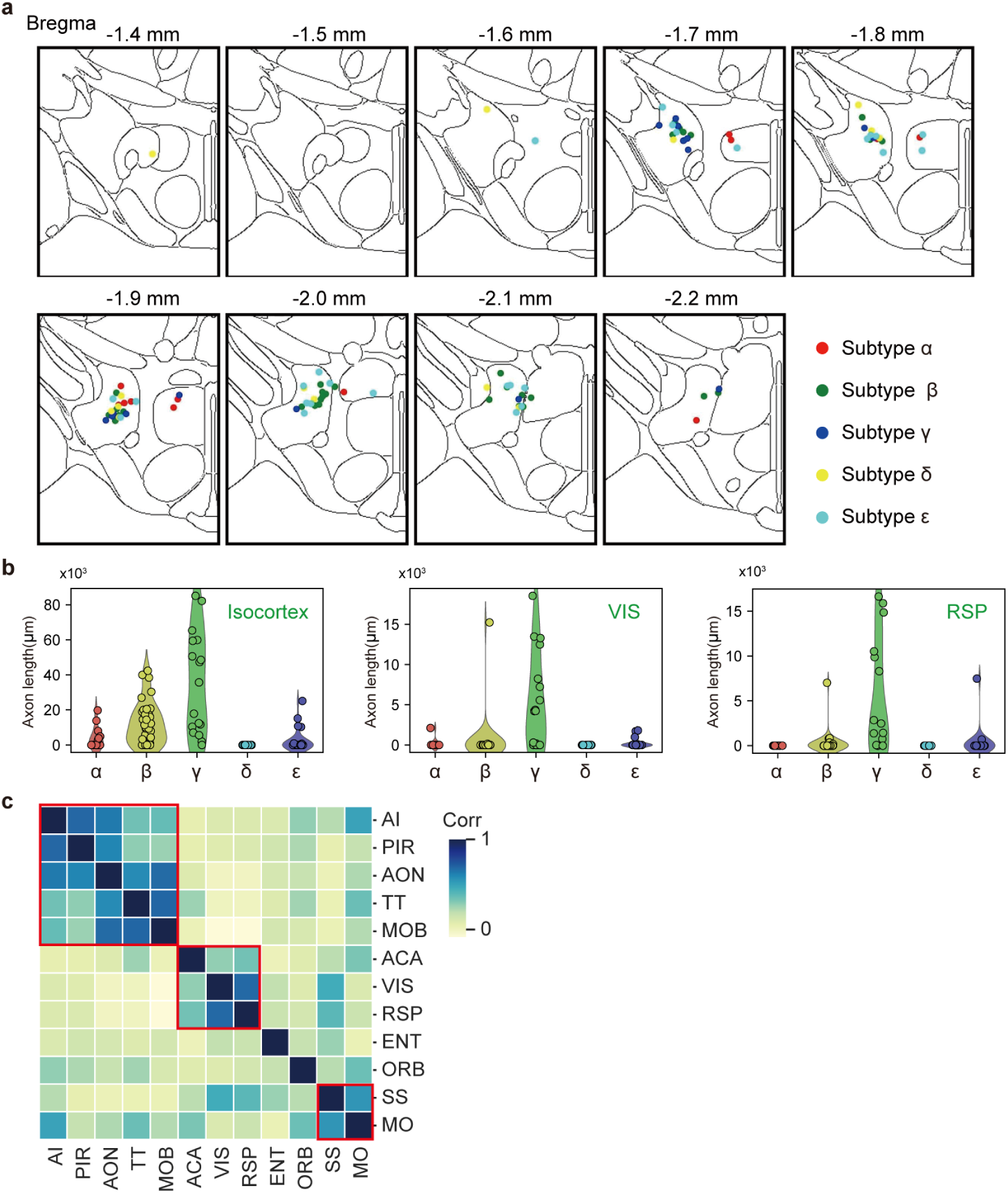
Projectome-defined *Orexin* neuron subtypes, *related to Fig. 4.* **a.** The soma location of neurons in the five projectome-defined *Orexin* subtypes. **b.** Violin plots of individual neurons’ projection length (in μm) for the five projectome subtypes in specific cortical areas. Each circle represents an individual neuron. **c.** Correlation (corr) analysis of the projection length of individual *Orexin* neurons in different cortical domains identifies co-innervated areas highlighted in red boxes.

**Extended Data Fig. 8.**
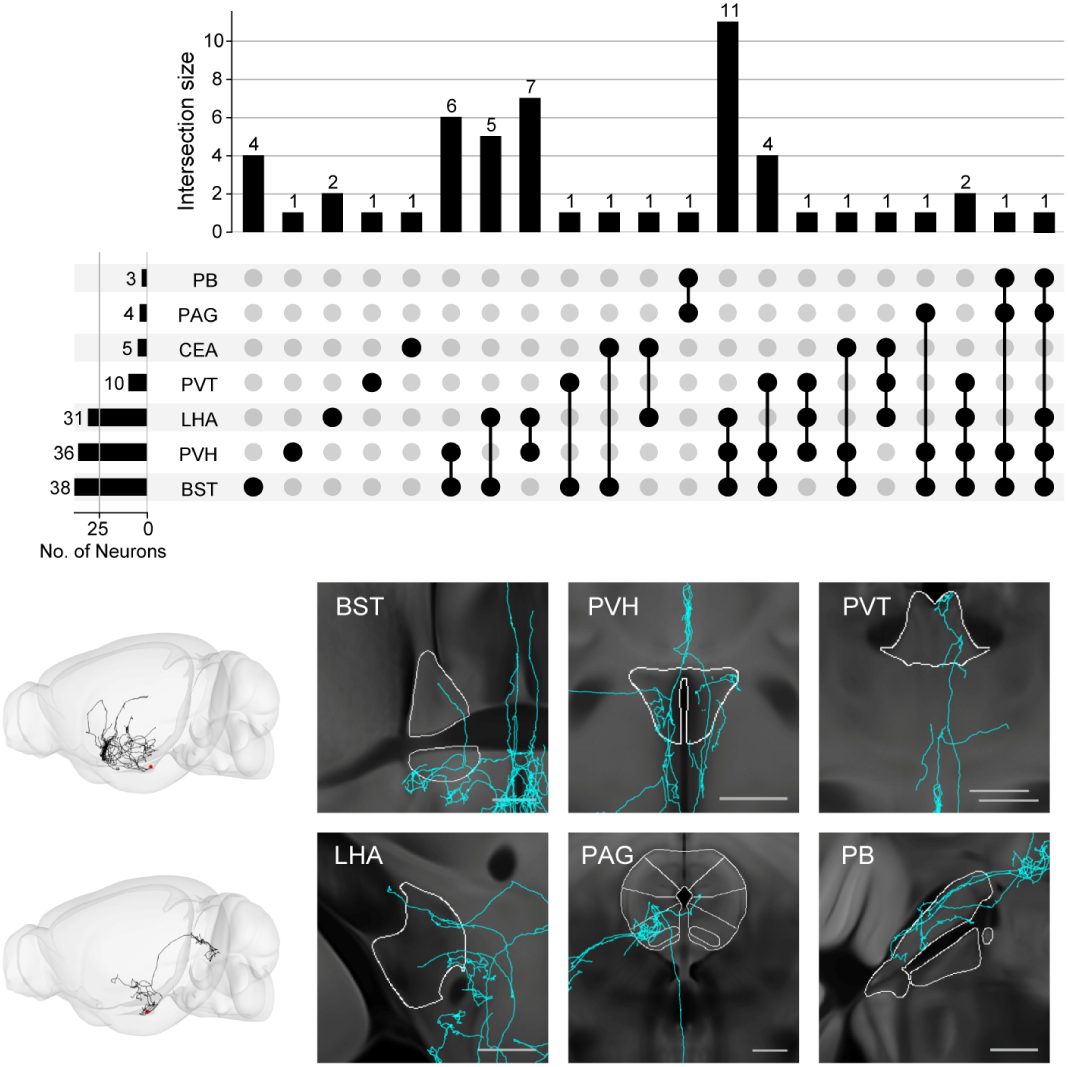
Axon collaterals of arcuate Agrp neurons, *related to Figure 5*. Top, An upset plot showing the intersection size of Agrp neurons that send axons to multiple brain areas indicated on the left. Bottom, axon projections of two representative Agrp neurons in 3D brain view on the left and the indicated target areas on the right. Both neurons projected to multiple targets.

## Supplementary Tables

**Supplementary Table 1.**
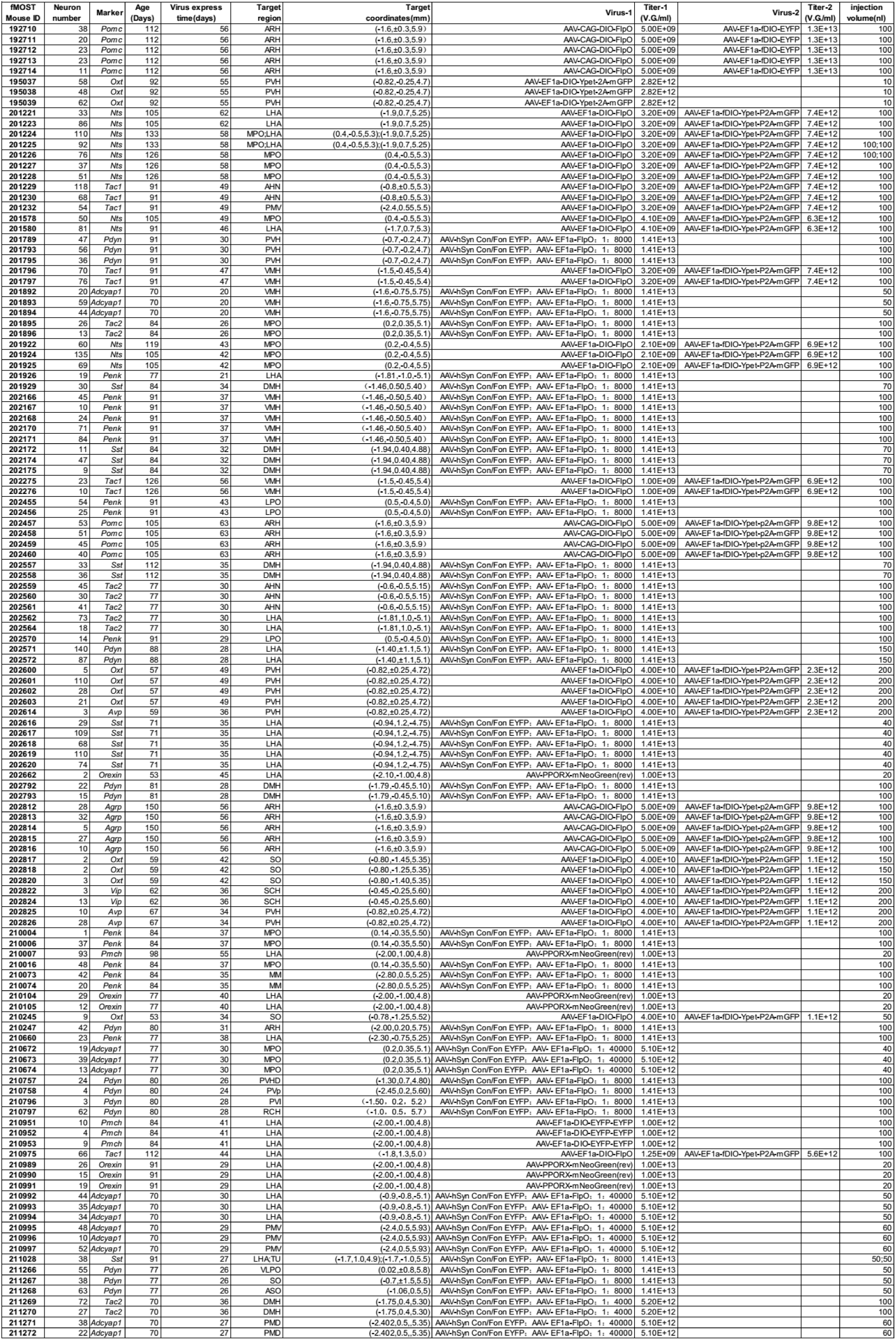
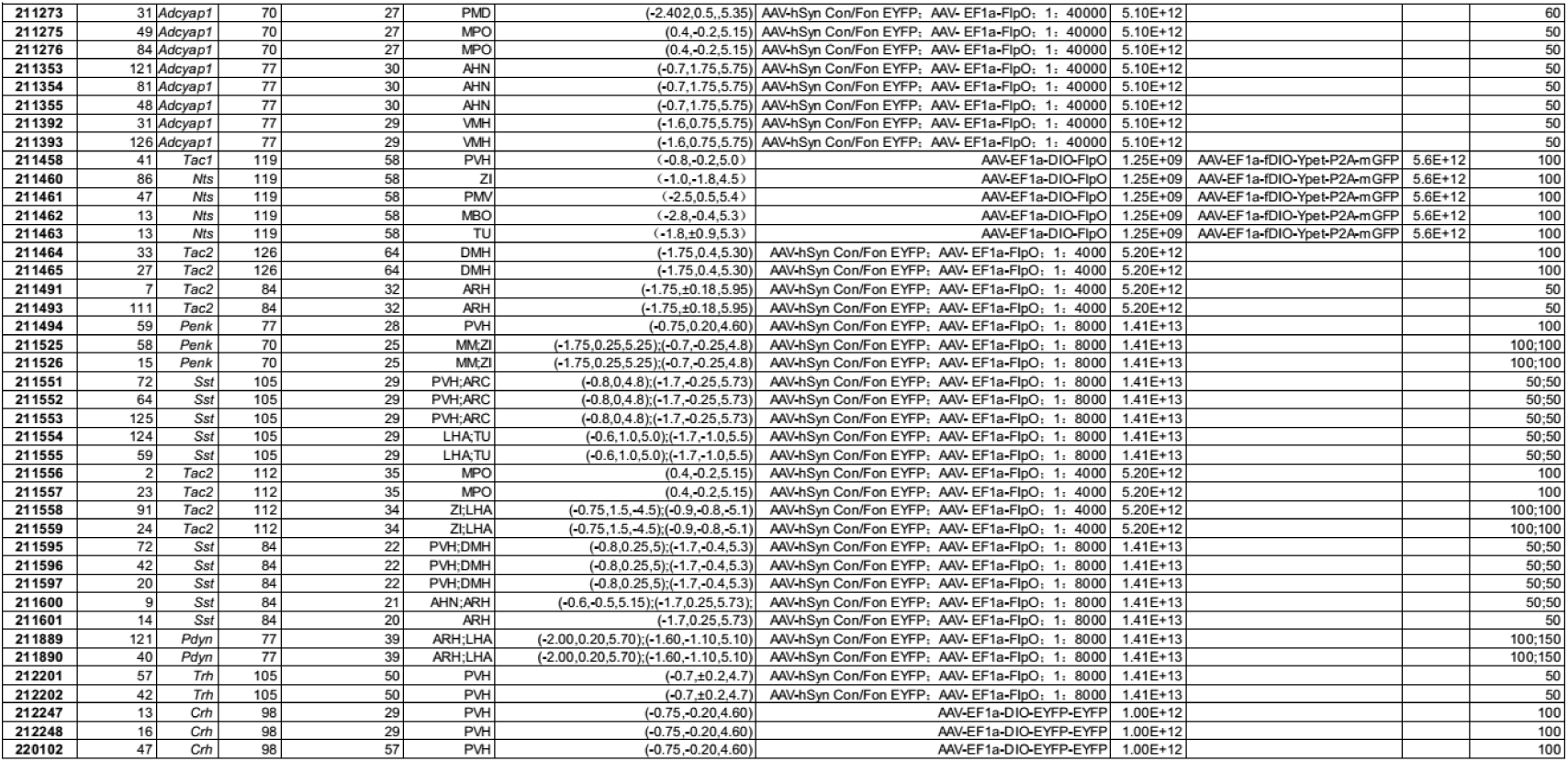
Information of the brain samples imaged, *related to Fig. 1*. In this table, we listed the age (in days) and genotype of the animals used for injection for each brain sample, and the volume and titer of viruses injected, and the injection coordinates. We also listed the number of neurons successfully reconstructed from each imaged sample. Note that all reconstructed neurons underwent several quality check rounds and only represent a fraction of neurons labeled in each brain sample.

**Supplementary Table 2.**
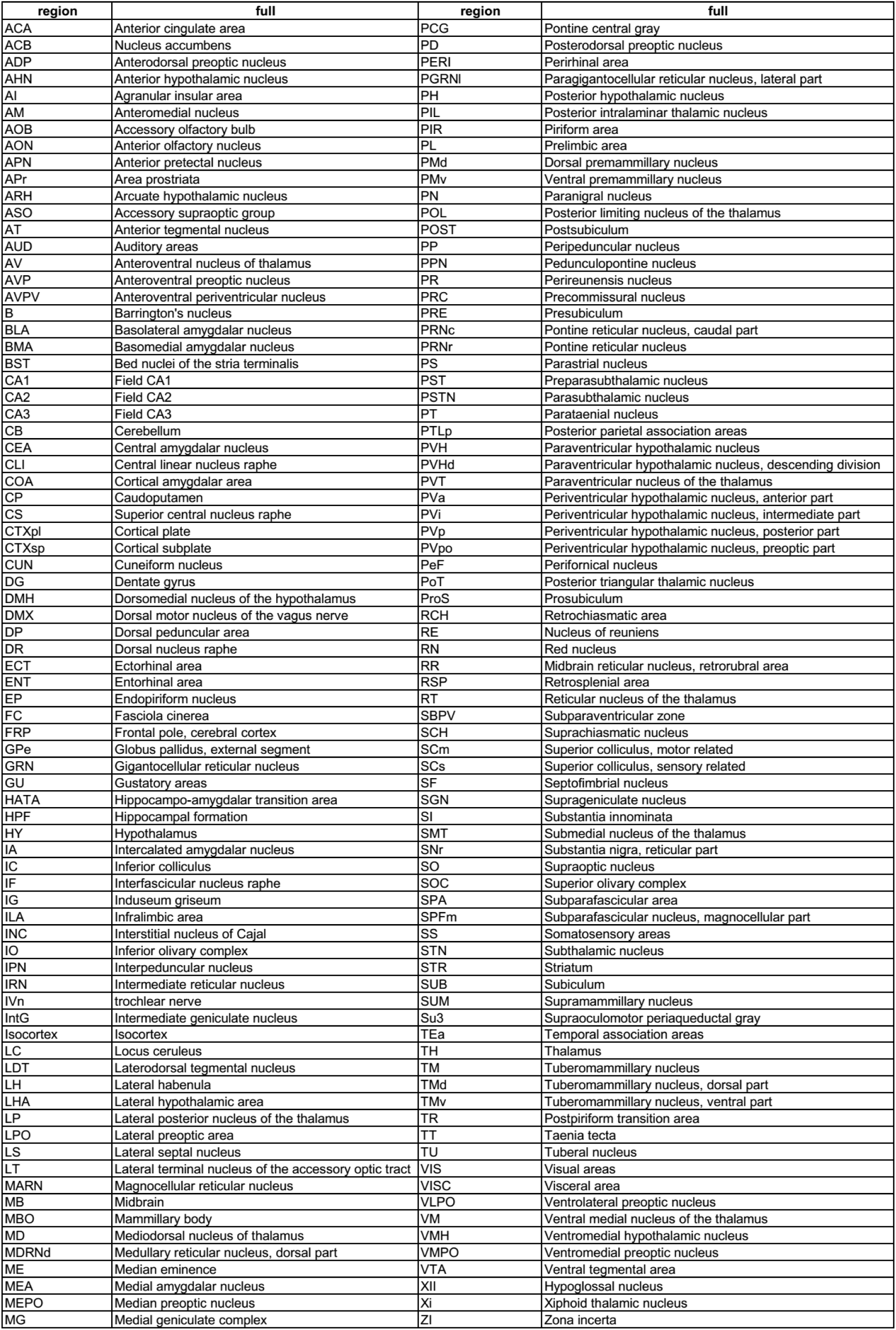
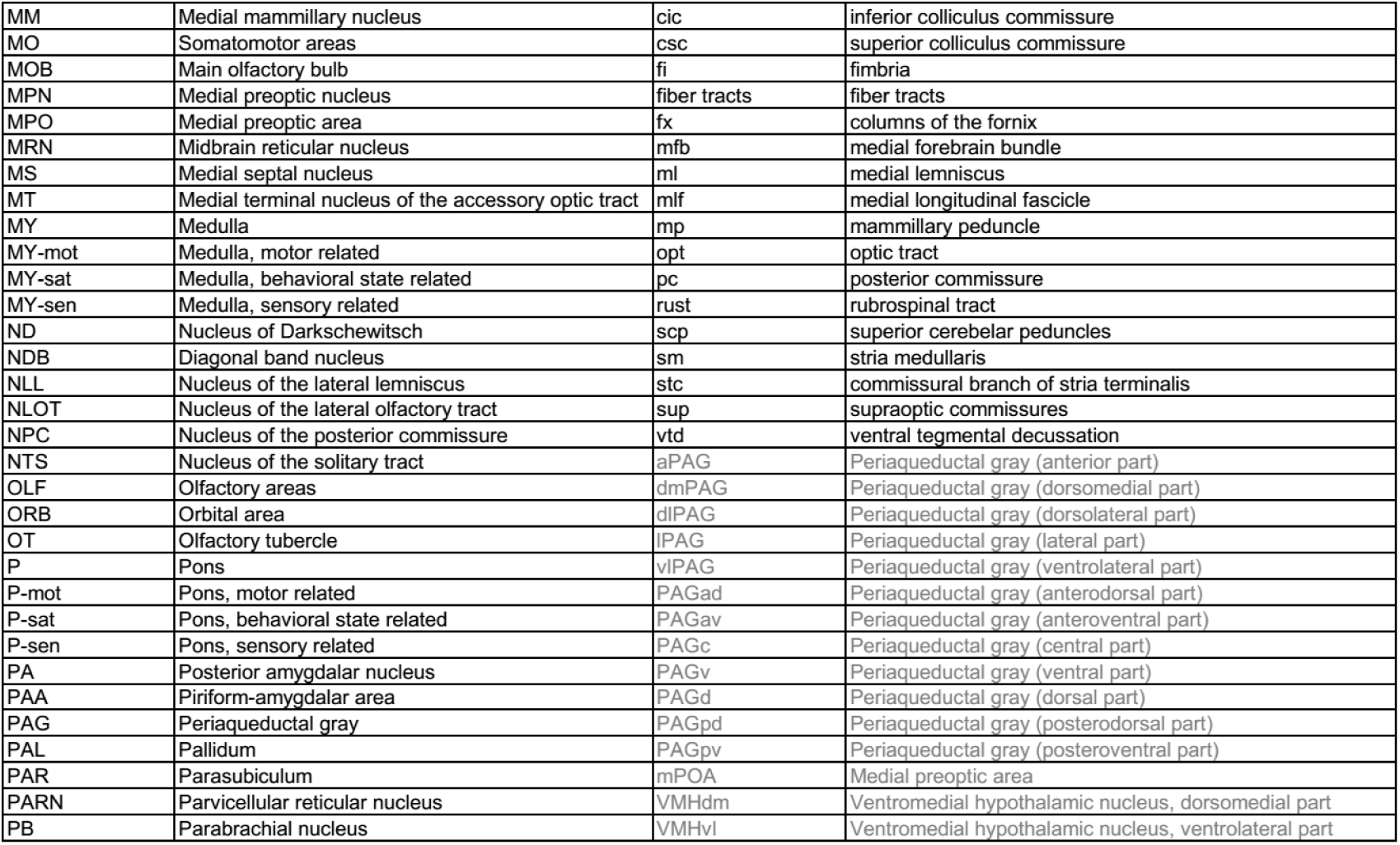
Nomenclature and Abbreviation of brain structures, *related to Fig. 1–7*. All listed brain areas’ names were consistent with the Allen CCFv3.0 except for colored ones, which were based on the Paxinos and Franklin Mouse Brain Atlas (2nd edition) or manual demarcation.

**Supplementary Table 3.** The number of neurons in each subtype (#1-#31) that expressed various neuropeptides, *related to Fig. 1–2*.

| Subtypes | Adcyap1 | Agrp | Avp | Crh | Nts | Oxt | Pdyn | Penk | Pmch | Pomc | Sst | Tac1 | Tac2 | Trh | Vip | Orexin |
| --- | --- | --- | --- | --- | --- | --- | --- | --- | --- | --- | --- | --- | --- | --- | --- | --- |
| 1 | 29 | 6 | 0 | 0 | 12 | 3 | 17 | 17 | 2 | 19 | 16 | 11 | 12 | 7 | 0 | 1 |
| 2 | 8 | 0 | 0 | 0 | 27 | 1 | 1 | 1 | 0 | 0 | 0 | 0 | 1 | 0 | 0 | 1 |
| 3 | 0 | 0 | 0 | 0 | 0 | 0 | 0 | 2 | 1 | 2 | 0 | 2 | 1 | 3 | 0 | 5 |
| 4 | 36 | 1 | 0 | 3 | 11 | 0 | 7 | 31 | 2 | 1 | 6 | 15 | 7 | 3 | 1 | 2 |
| 5 | 6 | 3 | 0 | 0 | 2 | 1 | 6 | 8 | 1 | 11 | 6 | 3 | 7 | 2 | 0 | 4 |
| 6 | 4 | 0 | 1 | 3 | 10 | 17 | 4 | 7 | 1 | 1 | 36 | 8 | 5 | 2 | 0 | 0 |
| 7 | 22 | 0 | 0 | 4 | 15 | 0 | 21 | 9 | 3 | 1 | 31 | 10 | 9 | 2 | 0 | 1 |
| 8 | 49 | 2 | 0 | 3 | 31 | 0 | 13 | 12 | 5 | 3 | 34 | 15 | 12 | 2 | 1 | 0 |
| 9 | 60 | 11 | 1 | 2 | 56 | 1 | 19 | 22 | 7 | 24 | 59 | 25 | 34 | 7 | 3 | 0 |
| 10 | 58 | 4 | 1 | 2 | 23 | 1 | 10 | 13 | 3 | 15 | 45 | 20 | 11 | 4 | 3 | 4 |
| 11 | 33 | 20 | 0 | 1 | 23 | 1 | 7 | 18 | 2 | 18 | 42 | 9 | 17 | 10 | 0 | 0 |
| 12 | 73 | 18 | 1 | 1 | 59 | 3 | 63 | 47 | 10 | 40 | 125 | 24 | 30 | 17 | 1 | 2 |
| 13 | 6 | 0 | 23 | 23 | 5 | 180 | 29 | 1 | 0 | 1 | 8 | 23 | 0 | 7 | 0 | 0 |
| 14 | 22 | 17 | 0 | 2 | 27 | 5 | 32 | 30 | 5 | 26 | 52 | 12 | 9 | 0 | 0 | 0 |
| 15 | 34 | 6 | 0 | 2 | 31 | 5 | 47 | 24 | 6 | 21 | 76 | 13 | 24 | 5 | 1 | 0 |
| 16 | 16 | 0 | 3 | 4 | 14 | 68 | 32 | 10 | 2 | 2 | 17 | 3 | 4 | 1 | 0 | 9 |
| 17 | 21 | 0 | 0 | 5 | 14 | 17 | 55 | 18 | 3 | 5 | 14 | 5 | 12 | 2 | 0 | 2 |
| 18 | 5 | 0 | 0 | 0 | 2 | 2 | 11 | 4 | 1 | 3 | 3 | 3 | 2 | 0 | 0 | 37 |
| 19 | 8 | 0 | 0 | 1 | 23 | 1 | 13 | 7 | 2 | 7 | 14 | 1 | 35 | 1 | 0 | 0 |
| 20 | 7 | 0 | 2 | 6 | 0 | 25 | 16 | 1 | 1 | 0 | 0 | 0 | 0 | 0 | 0 | 4 |
| 21 | 33 | 0 | 2 | 0 | 95 | 5 | 29 | 42 | 6 | 4 | 73 | 18 | 54 | 0 | 0 | 8 |
| 22 | 32 | 0 | 0 | 0 | 20 | 0 | 13 | 9 | 0 | 0 | 0 | 3 | 24 | 0 | 0 | 0 |
| 23 | 34 | 7 | 0 | 1 | 35 | 0 | 13 | 18 | 0 | 14 | 42 | 7 | 36 | 0 | 0 | 5 |
| 24 | 22 | 0 | 0 | 2 | 25 | 1 | 21 | 35 | 2 | 6 | 54 | 5 | 50 | 0 | 0 | 2 |
| 25 | 42 | 0 | 0 | 2 | 26 | 1 | 127 | 34 | 2 | 19 | 48 | 93 | 32 | 0 | 0 | 2 |
| 26 | 57 | 1 | 0 | 2 | 32 | 0 | 24 | 34 | 2 | 1 | 93 | 18 | 38 | 0 | 0 | 5 |
| 27 | 17 | 0 | 0 | 0 | 7 | 0 | 21 | 6 | 1 | 8 | 8 | 12 | 16 | 0 | 0 | 0 |
| 28 | 129 | 3 | 4 | 5 | 142 | 3 | 86 | 45 | 19 | 16 | 114 | 88 | 58 | 14 | 2 | 0 |
| 29 | 56 | 1 | 1 | 0 | 43 | 2 | 57 | 26 | 7 | 24 | 33 | 25 | 22 | 3 | 0 | 2 |
| 30 | 73 | 0 | 1 | 0 | 103 | 3 | 16 | 59 | 10 | 2 | 66 | 23 | 27 | 4 | 4 | 6 |
| 31 | 56 | 2 | 1 | 2 | 126 | 2 | 45 | 59 | 10 | 10 | 80 | 32 | 74 | 3 | 0 | 1 |

**Supplementary Table 4.**
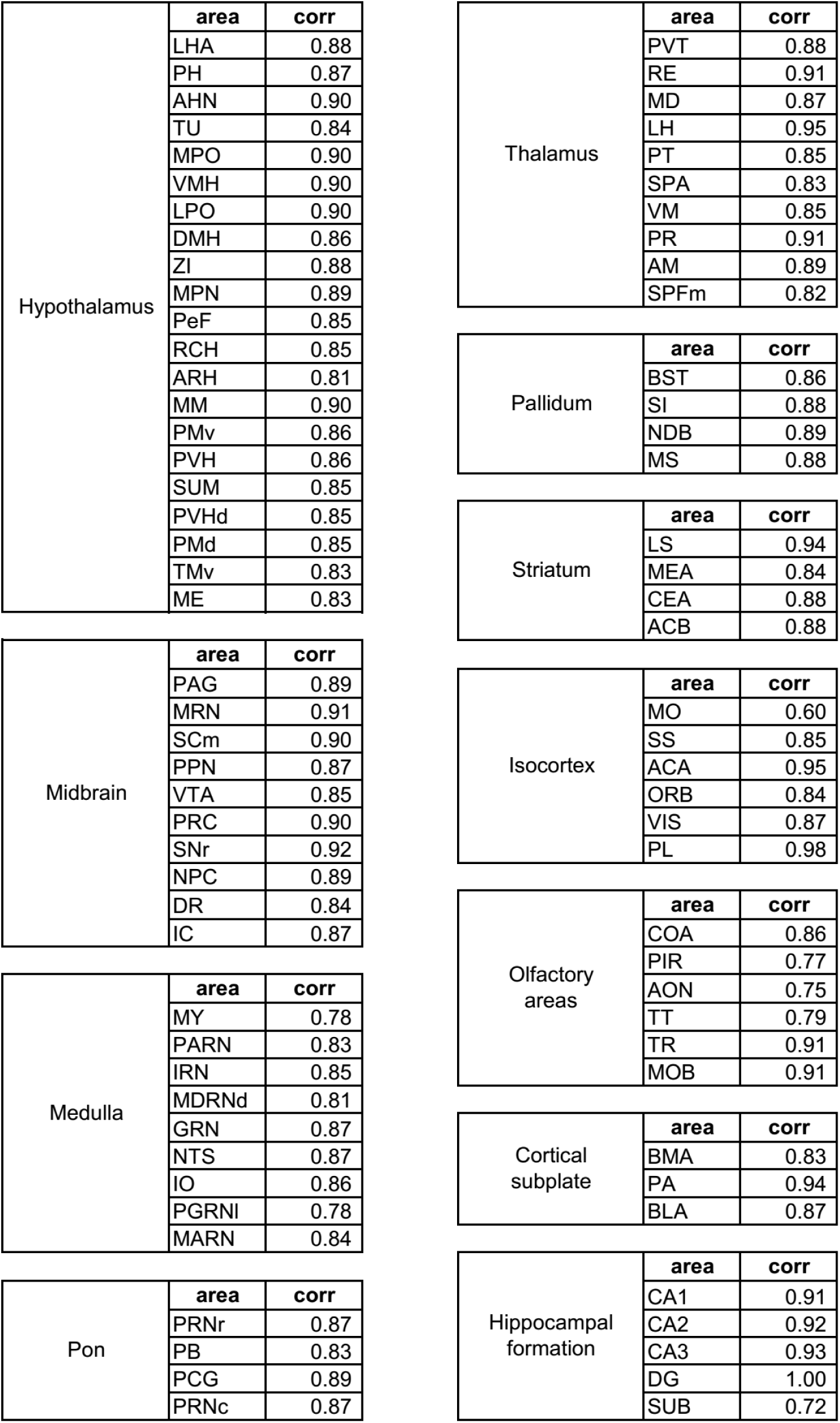
Correlation co-efficiency of axon projection length and synaptic terminal counts in each brain region. Correlation in all brain regions were highly significant with the *p* value approaching 0.

